# Computational biology predicts metabolic engineering targets for increased production of 102 valuable chemicals in yeast

**DOI:** 10.1101/2023.01.31.526512

**Authors:** Iván Domenzain, Yao Lu, Junling Shi, Hongzhong Lu, Jens Nielsen

## Abstract

Development of efficient cell factories that can compete with traditional chemical production processes is complex and generally driven by case-specific strategies, based on the product and microbial host of interest. Despite major advancements in the field of metabolic modelling in recent years, prediction of genetic modifications for increased production remains challenging. Here we present a computational pipeline that leverages the concept of protein limitations in metabolism for prediction of optimal combinations of gene engineering targets for enhanced chemical bioproduction. We used our pipeline for prediction of engineering targets for 102 different chemicals using *Saccharomyces cerevisiae* as a host. Furthermore, we identified sets of gene targets predicted for groups of multiple chemicals, suggesting the possibility of rational model-driven design of platform strains for diversified chemical production.

**One sentence summary:** Novel strain design algorithm ecFactory on top of enzyme-constrained models provides unprecedented chances for rational strain design and development.

## Introduction

The accelerated rise of metabolic engineering, the rewiring of cells metabolism for enhanced production of metabolites^1^, and synthetic biology, the assemble of novel synthetic biological components and their integration into cells^2^, has enabled the development of microbial strains with increased production capabilities of chemicals from renewable feedstocks. These engineered microbes, also known as microbial cell factories (MCF), have been generated for production of multiple specialized compounds, such as pharmaceuticals^3,4^, biofuels^5,6^, food additives^7,8^ and platform chemicals^9^. Most of these cases have relied on use of the bacterium *Escherichia coli* and the yeast *Saccharomyces cerevisiae* as platform cell factories. Despite success in development of many processes, complete development of MCFs usually takes several years of research and costs USD50M, on average, in order to bring a proof-of-concept strain forward for commercial production^10^.

As metabolism is a complex and highly interconnected network, the time and resource intensive process of MCF development can be alleviated by the use of genome-scale metabolic models (GEMs) together with computational algorithms, aiming to find non-intuitive gene engineering targets for enhanced production^11^. Several methods for MCF design have been developed in past years and used to drive metabolic engineering projects such as production of lycopene^12,13^ and lactate^14^ in *E. coli*, and drug precursors in *S. cerevisiae* cells^15^. However, the most widely used methods for MCF design (MOMA^16^, FSEOF^12^, optKnock^17^ and optForce^18^) tend to predict extensive lists of gene target candidates, and modelers often find themselves in need of imposing custom criteria to delimit the number of candidate gene targets to be tested, in order to reduce the amount of experimental work. Additionally, state-of-the-art GEMs tend to overpredict metabolic capabilities of cells due to the lack of kinetic and regulatory information in their formulation, hindering their applicability for further quantitative evaluation and comparison of predicted metabolic engineering strategies. Kinetic models have also been used for the development of strain design algorithms, such as k-OptForce^19^, however, the limited size of this kind of models impedes prediction of metabolic gene targets in a genome-scale^20^.

Here we present a computational method (ecFactory) for prediction of optimal metabolic engineering strategies, that circumvents the problem of arbitrary selection of the number of gene candidates by leveraging the vast amount of enzymatic capacity data, together with the improved phenotype prediction capabilities, of enzyme-constrained metabolic models (ecModels, generated by the GECKO toolbox)^21^. The performance of ecFactory was systematically tested and evaluated by comparing the predictions with experimental data for multiple study cases. Using this method we identified gene targets for increased production of 102 different chemicals in *S. cerevisiae*, enabling identification of gene targets common to multiple groups of products, suggesting the opportunity for development of platform strains that can be used for diverse chemical production. Moreover, our analysis quantitative estimation of enzyme- and substrate-limitations for production of the 102 studied chemical products. To enable wider utilization of these results by the community, we established a web-based resource for accessible query and visualization of the gene target predictions in the context of Metabolic Atlas, and we expect this resource to facilitate significant advancements in development of yeast MCFs through metabolic engineering.

## Results and discussion

### Modelling production of 102 chemical products in yeast

A list of 102 industrially relevant natural products, whose metabolic production pathways are known and reported in the literature, was collected. Products were grouped into 10 different families according to their chemical characteristics: amino acids (26), terpenes (22), organic acids (15), aromatic compounds (9), fatty acids and lipids (9), alcohols (8), alkaloids (6), flavonoids (5), bioamines (2) and stillbenoids (1). From these, 50 products were found to be native metabolites in *S. cerevisiae*, whilst 52 products were identified as heterologous, according to an enzyme-constrained metabolic model for yeast (ecYeastGEM v8.3.4)^21^. A summary of the chemical classification of products is shown in **Fig. 1A** and **supp. table S1**. Production pathways were reconstructed for all these heterologous products and incorporated into ecYeastGEM, taking energy and redox requirements as well as reported kinetic data into account (see **Materials and Methods**). All of the 53 reconstructed heterologous pathways are described in **supp. table S2**.

**Figure 1.**
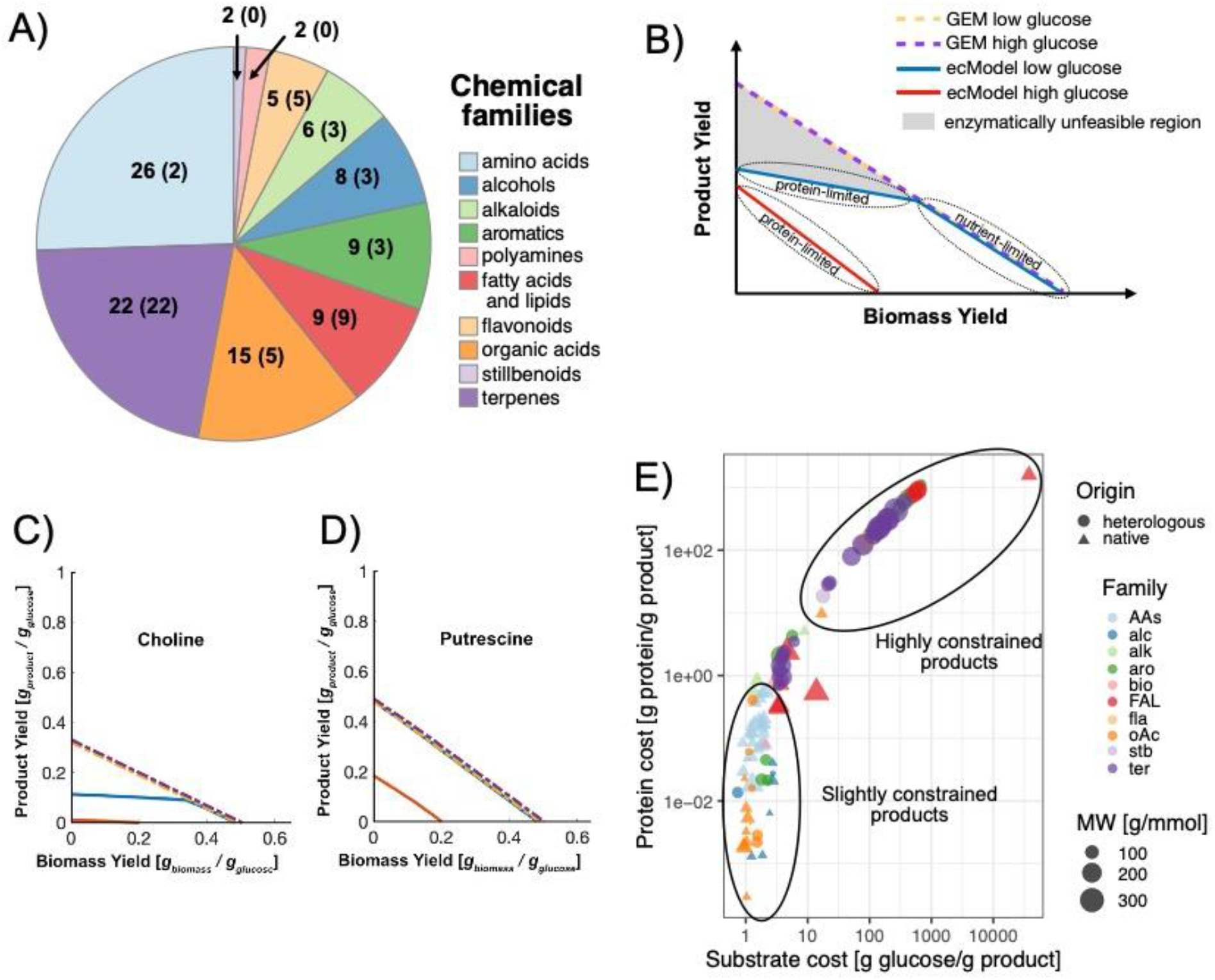
Exploration of chemical production in yeast using enzyme-constrained metabolic modeling. A) Chemical classification of 102 chemicals for in silico prediction. Numbers within parenthesis indicate number of native products in the different families, those outside the parenthesis indicate the total number of products in the family. B) Production landscape predicted by a metabolic model with and without enzyme constraints at low and high glucose uptake levels. C) Production yield plot for the highly protein-constrained product choline. D) Production yield plot for the substrate-limited putrescine. E) Predicted substrate and protein cost of chemical production in yeast. Product origin, chemical classification and molecular weights are indicated by the characteristics of the 102 markers.

### In silico assessment of production capabilities for 102 chemicals in yeast using metabolic modeling with enzyme constraints

The production capabilities of *S. cerevisiae* were quantitatively explored, using both YeastGEM and ecYeastGEM, by computing optimal production yields for all of the 102 studied chemicals, constrained by low and high glucose consumption regimes (1 mmol/gDw h; and 10 mmol/gDw h) and biomass production rates spanning the range between zero and a maximum attainable value, using flux balance analysis (FBA) simulations^22^.

As FBA relies on optimality principles, usually assuming maximization of cellular growth as a cellular objective^23^, there is a trade-off between biomass formation and accumulation or secretion of products of interest. Yeast has evolved the ability to switch to mixed respiro-fermentative metabolic regimes when nutrients are available in excess, favoring enzymatic efficiency over biomass yield on substrate^24–27^. As ecModels account for a limited enzymatic machinery in cells, different production capabilities are predicted when changing from low to high glucose uptake rate, in contrast to classic GEMs, that solely rely on stoichiometric constraints. This additional constraint results in a different production phase-plane as illustrated by **Fig. 1B**, i.e., instead of the standard linear trade-off between product formation and biomass formation there will be a regime where the product formation is limited by the protein constraint. Furthermore, the phase-plane becomes dependent on the glucose-uptake rate, such that at high glucose consumption the ecModel predicts a protein-limited regime of production, yielding lower production levels and biomass formation per unit of glucose. Protein limitations may also arise at low glucose consumption levels, for cases in which the production pathways for the chemical of interest involve inefficient enzymes (low specific activity). This introduces enzymatically unfeasible regions in the production space of a cell, indicated by the grey region in **Fig. 1B**. A typically protein-constrained production landscape with a region of difference between YeastGEM and ecYeastGEM predictions in the low glucose regime is shown for the alkaloid choline in **Fig. 1C**. In contrast, a production landscape solely governed by stoichiometric constraints at low glucose levels is shown in **Fig. 1D** for the polyamine putrescine. Additional examples of yield plots for chemicals belonging to all studied families can be found in **Fig. S1**.

Highly protein-constrained products were found by identifying those chemicals whose maximum production level demands the totality of the available enzyme mass in the model, at low levels of glucose consumption. In total, 40 out of the 53 analyzed heterologous products were found to be highly protein-constrained, in comparison to production of native metabolites, for which just 5 products were classified as part of the same group (**Fig. S2A**). Furthermore, strong protein limitations arise often for groups of heterologous chemicals derived from a native pathway with high enzymatic demands, such as terpenes and flavonoids, derived from the mevalonate pathway. On the other hand, few strongly protein-limited products were found amongst families connected to native biosynthetic processes, such as amino acids, organic acids and diverse alcohols (**Fig. S2B**). Protein constrained models offer the possibility of computing optimal costs of chemical production both in terms of substrate and required protein mass. Minimal protein and substrate mass costs per unit mass of product were computed for each of the 102 products (see **Materials and Methods** for further details), as has been previously suggested by other computational work^28^. **Fig. 1E** shows that a positive correlation between these two production costs exists, allowing the identification of slightly and highly constrained groups of products, with an overrepresentation of native products (amino acids, organic acids and some alcohols) in the former group, and heterologous chemicals (terpenes, flavonoids and some aromatic compounds) in the latter. This plot shows that for heterologous products, it is usually necessary to invest on improving enzyme properties, i.e., increase their catalytic efficiency, whereas for native products it is predominantly stoichiometric constraints that should be considered for minimizing costs. Moreover, it was found that slightly constrained products tend to be lighter, in terms of molecular weight, than those in the highly constrained group. This is also suggested by substrate costs, as larger organic molecules require more carbon to be formed, notwithstanding, this also suggests that a heavier enzymatic burden is needed for assembly of large molecules, as it is likely that additional, and less efficient, enzymatic steps are involved in their synthesis.

The effect of increasing enzyme catalytic efficiency for improving production levels was explored with FBA simulations with ecYeastGEM at different activity levels of rate-limiting enzymes. For highly protein-constrained products, such as the alkaloid psilocybin, a monotonic linear decrease of the substrate cost is observed when decreasing the total production protein cost by enhancing the activity of the heterologous tryptamine 4-monooxygenase (P0DPA7). **Fig. S3A** shows that when the P0DPA7 catalytic efficiency is increased by 100-fold, the total oxygen consumption is predicted to increase by 75%, which suggests that reducing the protein burden of the psilocybin biosynthetic pathway releases protein mass that can be used by the cell to meet its energy demands by an increased respiratory rate. Overall, this metabolic rewiring shifts the psilocybin production space in a direction of higher product yields (**Fig. S3B**). However, the product yield is still low indicating that other enzymes in the pathway may have to be improved to further increase yield. A similar behavior was obtained for the case of valencene, a moderately protein-constrained terpenoid, by increasing activity levels of the sole heterologous limiting enzyme, terpene synthase (S4SC87), from 1 to a 100-fold. A positive correlation was also observed between substrate and protein costs for this product (**Fig. S3C**), however, lower slopes in the production cost space were obtained for higher activity values of S4SC87. **Fig. S3D** illustrates that increased activity of this limiting enzyme reduces the enzymatically unfeasible region of the valencene production space, bringing its optimal production line closer to the stoichiometrically constrained limit (blue and dark red lines).

In sum, model predictions indicate that heavily protein-constrained biosynthetic pathways could result in the increase of protein and substrate costs of production. This kind of pathways require resources from the limited cellular enzymatic machinery, hence, the substrate-efficient respiratory pathway for energy production is compromised in favor of substrate-inefficient fermentative pathways, which reduces the protein burden necessary for sustaining cellular energy levels.

### An integrative constraint-based method for prediction of metabolic engineering strategies

The flux scanning with enforced objective function algorithm (FSEOF)^12^ has been extensively used for identification of metabolic engineering targets in yeast, due to its implicit consideration of the tradeoff between biomass and metabolite production. It is of particular interest to explore this method in the context of ecModels as variable energetic and biosynthetic requirements may induce a complete change of the cellular behavior. Therefore, engineering strategies that minimize the substrate and protein costs for optimal bio production can be predicted, furthermore, predictions have boosted heme accumulation in yeast cells by 70-fold^29^. In order to ensure predictive robustness and minimizing the number of false positives among predictions, we revised and systematized this approach and developed ***ecFactory***, a multi-step constrained-based method for prediction of engineering gene targets for enhanced biochemical production, based on the principles of FSEOF and on the ability of ecModels to compute enzyme demands for biochemical reaction, providing systematic criteria to predict an optimal minimal set of modifications for increasing production of target metabolites.

In summary, ecFactory consists of three basic steps: 1) prediction of gene expression scores, indicating intensity and directionality of genetic modifications; 2) discard gene targets encoding for unfavorable enzymes (redundant, low efficiency) and; 3) Obtention of a minimal combination of modifications required for driving cells from optimal biomass formation to a metabolic production regime. The overall objective of this method is to obtain a reduced list of gene targets, focusing on the optimal strategies for enhanced production by taking enzyme allocation and connectivity into account. All the constituent steps of the ecFactory method are illustrated in **Fig. 2** and explained in detail in the **Materials and Methods** section of the **Supplementary Materials**.

**Figure 2.**
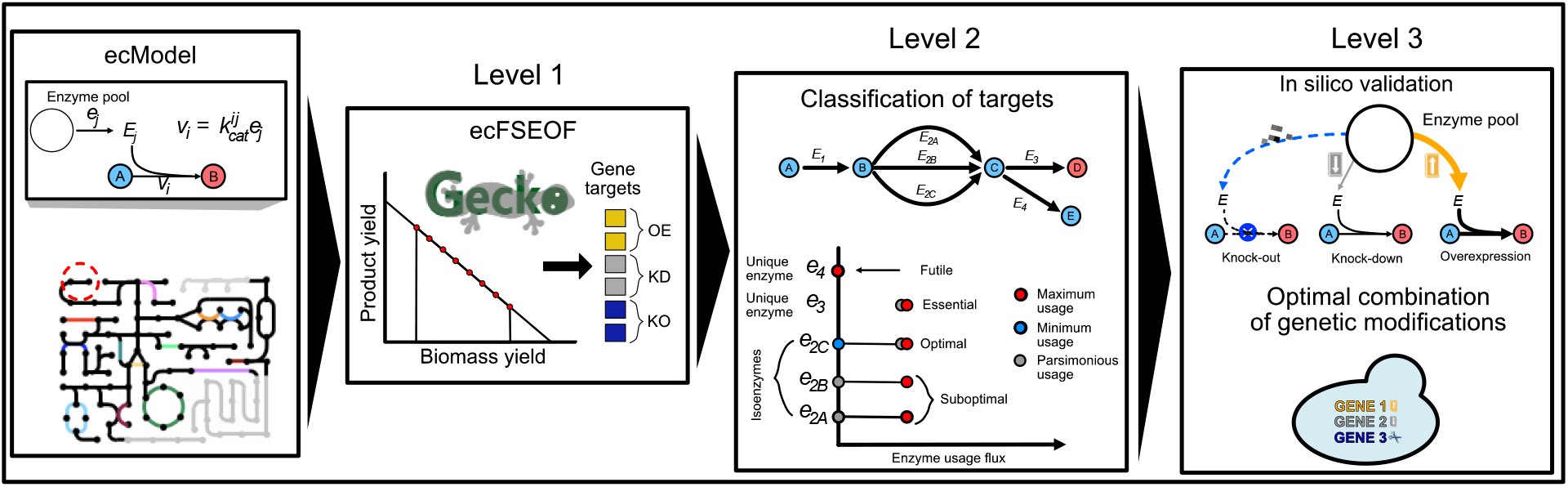
Prediction of metabolic engineering targets with ecFactory. A metabolic model with enzyme constraints is used for (1) prediction of gene targets for rewiring flux towards increased production. (2) Gene targets are classified and filtered according to enzymatic efficiency and connectivity. (3) A minimal combination of targets for sustaining optimal production levels is obtained.

Furthermore, the classification of targets according to the characteristics of their respective enzymes (illustrated by **Fig. S4**), facilitates a deeper understanding of the predicted optimal metabolic engineering strategies. The list of 12 gene targets for 2-phenylethanol (**Table S4**) suggests that, in order to increase production of this chemical, enzymes that are optimal for providing the necessary metabolic precursors and cofactors are predicted as targets for overexpression. Knock-down and knock-out targets aim to direct the metabolic flux towards optimal production while reducing the formation of biomass precursors in excess (glycerolipids in this case).

### Enzyme constraints enable identification of optimal combinations of genetic modifications for 102 chemicals in yeast

The ecFactory method was used to predict gene targets for enhanced production of each of the 102 chemicals. The method proved to be effective at returning predictions for all cases, while reducing the number of candidate gene targets at each of its sequential steps. The distributions of the predicted number of gene targets per product (shown in **Fig. 3A)** shows the major contribution of classifying targets according to their enzymatic characteristics (step 2) at reducing the number of predicted OEs, KDs and KOs. On average the first step of the method (FSEOF), running on ecYeastGEM, predicted 85 gene targets per product (28 OEs, 42 KDs and15 KOs), the number of targets is then reduced by the following steps by 73%, as only optimal candidates are returned by the ecFactory algorithm (7 OEs, 9 KDs and 5 KDs per product, on average). Notably, predictions reveal that increasing production of protein-limited and heterologous chemicals require significantly more genetic modifications, compared to substrate-limited and native products (p-values = 1.16×10^-5^ and 2.3×10^-3^, respectively, under a one-sided two-sample Kolmogorov-Smirnov test) as shown in **Fig. S5**. These differences are caused by the large number of gene knock-downs and knock-outs that are required to change the energy production strategy from cellular respiration to a fermentative metabolism, so that the limited cellular enzyme capacity can be optimally allocated to the final production reaction steps, which tend to be inefficient for these kinds of products. A more detailed presentation of results, by chemical family, method steps, and target types, is available in **supp. table S3**.

**Figure 3.**
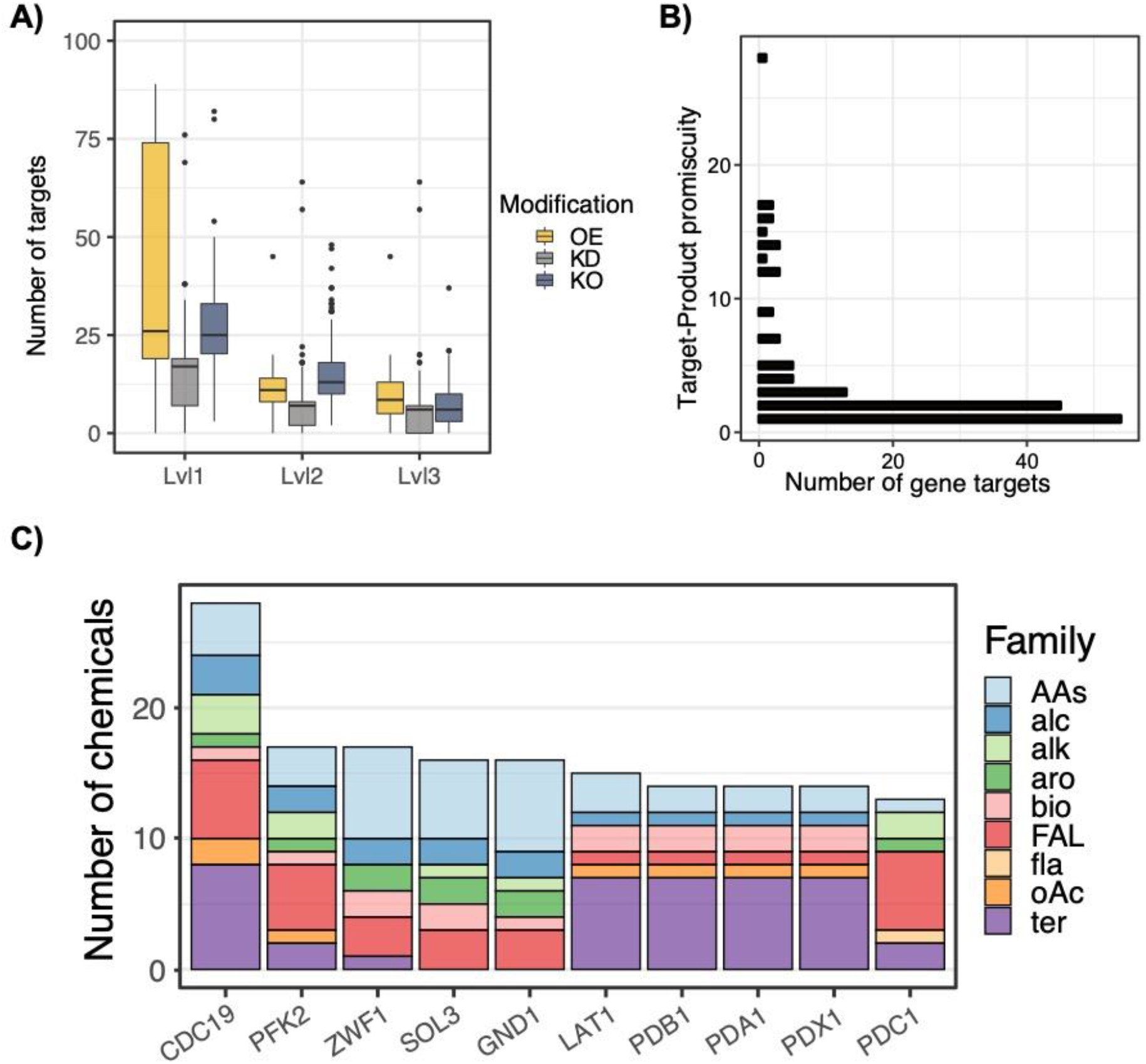
Prediction of gene engineering targets for increased production of 102 chemicals in yeast. A) Distribution of the number of gene targets per product predicted at different steps in the ecFactory pipeline. Level1, FSEOF-based prediction; Level2, filtering by enzyme characteristics; Level3, obtention of minimal set of targets for optimal production. B) Distribution of product specificity of gene targets across 102 chemicals. C) Representation of the presence of the top 10 most common predicted overexpression targets across products and families.

Overall, 150 endogenous genes in yeast are predicted as OE target for at least one of the modeled products; 88 different genes are predicted as KD targets and 129 as KO targets. More than 50% of the targets predicted for OE, KD and KO are specific to one or two of the 102 products (Fig.3B, Fig. S6A and Fig. S6C). Nonetheless, small sets of genes are predicted as targets for a high number of products (promiscuous targets), spanning almost all chemical classifications in this study. Genes encoding for reaction steps in the pentose-phosphate pathway and pyruvate metabolism, together with PFK2 in the glycolysis pathway, are predicted as the most common OE targets across products; the most common KD and KO gene targets encode for enzymes in the TCA cycle, oxidative phosphorylation and synthesis of biomass precursors (steroids, glycerolipids, nucleotides and amino acids), as shown in **Fig. 3C**, **Fig. S6B** and **Fig. S6D**, suggesting a global strategy of redirecting carbon flux into heterologous pathways and alternative energy production mechanisms.

### In silico predictions capture successful metabolic engineering strategies in yeast

It was found that 7 out of the 12 predicted gene targets to increase 2-phenylethanol have been previously engineered in *S. cerevisiae, Yarrowia lipolytica and Kluyveromyces marxianus* strains with enhanced 2-phenylethanol production levels^30–32^ (**Table S4**), indicating that ecFactory predictions can be capable of capture targets proposed by rational engineering approaches. As another example of experimentally validated predictions, the case of spermine is of particular interest. In this case, the ecFactory method was able to capture 9 of the implemented targets (MAT, ODC, SPE2, SPDS, MEU1 APT2 and PRS for OE; FMS1 and CAR2 for KO) in a successfully engineered strain for spermidine production, an immediate precursor of spermine^33^. It was also found that the experimental implementation of a heterologous cytosolic ornithine cycle was resembled by a general predicted overexpression of its native mitochondrial version.

These particular results suggest that the method is able to capture the underlying logic of highly complex rational engineering approaches that require the coordination of multiple sectors of metabolism, as shown by **Fig. S7**. Overall, the predicted gene modifications aim to increase spermine biosynthesis by overexpression of the whole ornithine cycle, a direct precursor, together with the Yang cycle and some steps in the pentose phosphate pathway (PPP) in order to increase S-adenosyl-L-methionine, another important precursor of polyamines. Interestingly, when focusing on the final predictions for this product (targets in step 3), just 5 of the 8 aforementioned genes were classified as optimal targets for spermine production (SPDS, ARG8 andARG5,6 OEs, together with FMS1 and CAR2 KO). This suggests that, according to enzyme capacity and metabolic connectivity, it is possible to reduce complex rational metabolic engineering strategies, to fewer modifications on crucial reaction steps in pathways that need to be rewired and coordinated, one of the purposes for which this method was designed.

In order to validate the quality of the ecFactory predictions, we searched the literature for independent experimental studies in *S. cerevisiae* that have been successful at increasing production levels of chemicals included in our list. Gene modifications validated for diverse chemicals were found to be predicted as optimal gene targets by ecFactory, shown in **Table 1**. Interestingly, several of these targets are common to multiple products. In total, 28 predicted different gene targets were found as experimentally validated across 22 products, which are also part of different chemical classes. The most repeated genes among these targets correspond to overexpression in the ergosterol, mevalonate, shikimate and polyamine biosynthesis pathways.

**Table 1.**
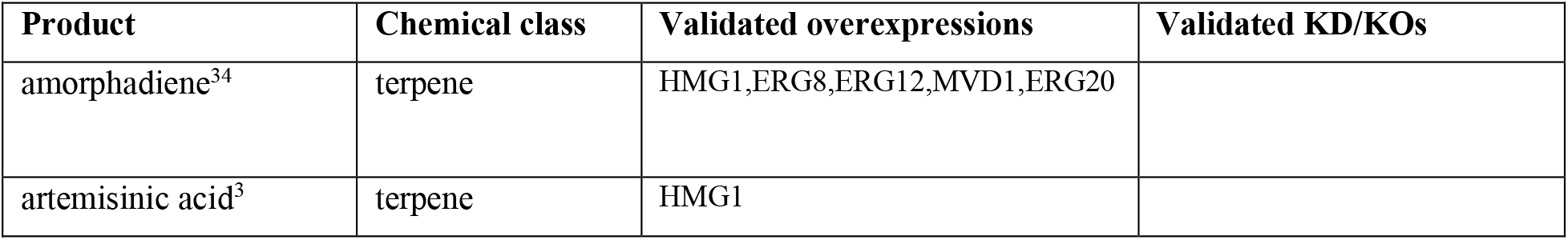
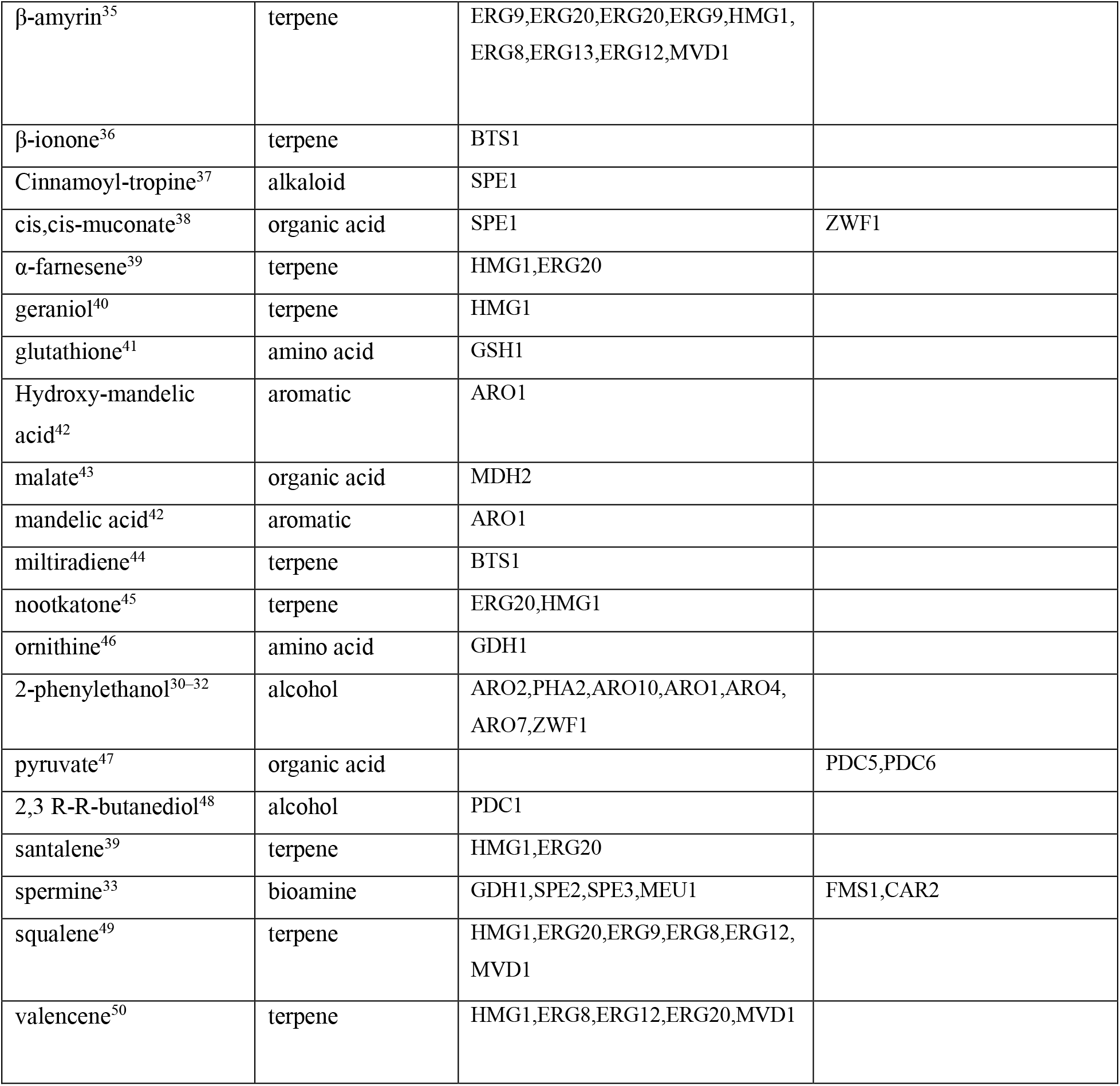
Predicted gene targets with experimental validation.

| Product | Chemical class | Validated overexpressions | Validated KD/KOs |
| --- | --- | --- | --- |
| amorphadiene<sup>34</sup> | terpene | HMG1,ERG8,ERG12,MVD1,ERG20 |  |
| artemisinic acid<sup>3</sup> | terpene | HMG1 |  |
| β-amyrin<sup>35</sup> | terpene | ERG9,ERG20,ERG20,ERG9,HMG1,ERG8,ERG13,ERG12,MVD1 | |
| β-ionone<sup>36</sup> | terpene | BTS1 | |
| Cinnamoyl-tropine<sup>37</sup> | alkaloid | SPE1 |  |
| cis,cis-muconate<sup>38</sup> | organic acid | SPE1 | ZWF1 |
| α-farnesene<sup>39</sup> | terpene | HMG1,ERG20 | |
| geraniol<sup>40</sup> | terpene | HMG1 |  |
| glutathione<sup>41</sup> | amino acid | GSH1 |  |
| Hydroxy-mandelic acid<sup>42</sup> | aromatic | ARO1 |  |
| malate<sup>43</sup> | organic acid | MDH2 |  |
| mandelic acid<sup>42</sup> | aromatic | ARO1 |  |
| multiradiene<sup>44</sup> | terpene | BTS1 |  |
| nootkatone<sup>45</sup> | terpene | ERG20,HMG1 |  |
| ornithine<sup>46</sup> | amino acid | GDH1 |  |
| 2-phenylethanol<sup>30-32</sup> | alcohol | ARO2,PHA2,ARO10,ARO1,ARO4,ARO7,ZWF1 |  |
| pyruvate<sup>47</sup> | organic acid |  | PDC5,PDC6 |
| 2,3 R-R-butanediol<sup>48</sup> | alcohol | PDC1 |  |
| santalene<sup>39</sup> | terpene | HMG1,ERG20 |  |
| spermine<sup>33</sup> | bioamine | GDH1,SPE2,SPE3,MEU1 | FMS1,CAR2 |
| squalene<sup>49</sup> | terpene | HMG1,ERG20,ERG9,ERG8,ERG12,MVD1 |  |
| valencene<sup>50</sup> | terpene | HMG1,ERG8,ERG12,ERG20,MVD1 |  |

These similarities at the gene and pathway level among predictions for different chemical products, suggest the existence of metabolic engineering strategies capable of providing the necessary precursors for increasing production of groups of chemicals. This kind of strategies have been sought in experimental metabolic engineering, following rational approaches, and have proved to be successful for the development of platform yeast strains for production of different groups of molecules such as opioids^4^ and other alkaloids^51,52^, polyketides^53^ and terpenes^39,54^. Furthermore, cumulative combination of individual genetic modifications in a production strain is needed for achieving meaningful flux towards the desired chemical^29^, therefore, it is desirable to identify multiple gene targets, encompassing multiple metabolic pathways, that constitute the chassis for robust and diversified chemical production.

Gene targets common to all products in a given chemical family were sought for all cases in this study. The only chemical family with common predicted targets was found to be flavonoids, with 9 KDs (ADO1, ATP19, IDP1, LPD1, MAE1, MDH2 MET6, PPA2 and SAH1) and 7 KOS (CAR2, FAA, FAA4, FDH1, RNR1, RNR3 and RNR4). This combination of targets reveals an engineering strategy that decreases the TCA cycle and respiratory fluxes, the amount of carbon going towards acetyl-CoA and posterior fatty acid synthesis, synthesis of amino acids derived from 2-oxoglutarate and nucleotides biosynthesis. Altogether, this shows an optimal way of allocating carbon flux and the limited enzymatic machinery of yeast for the biosynthetic pathways producing catechin, genistein, kaempferol, naringenin and quercetin. Nevertheless, the impact of these modifications on other biological processes, such as regulatory networks, is not accounted for in the metabolic model and should be further assessed.

#### Model-driven design of platform strains for diverse chemical production

As highly promiscuous gene targets, for all kind of modifications, were found to be predicted for products present in most of the studied chemical families, other sets of targets common to groups of multiple products may exist among the ecFactory predictions. In order to systematize the analysis of gene target profiles across products, the 102 lists of targets were represented as mathematical vectors (see **Materials and Methods** section of the **Supplementary Materials** and **Figure 4A** for further details). Highly similar gene expression vectors were identified using the t-distributed stochastic neighbor embedding method (t-SNE), which is suited for visualization and identification of clusters in high dimensional datasets^55^. Two-dimensional representation of t-SNE results facilitated identification of 8 different clusters of target vectors, representing different groups of products. Product clusters are shown in **Figure 4B**. Notably, gene targets common to all products in a group were found for all clusters (**Table 2**).

**Figure 4.**
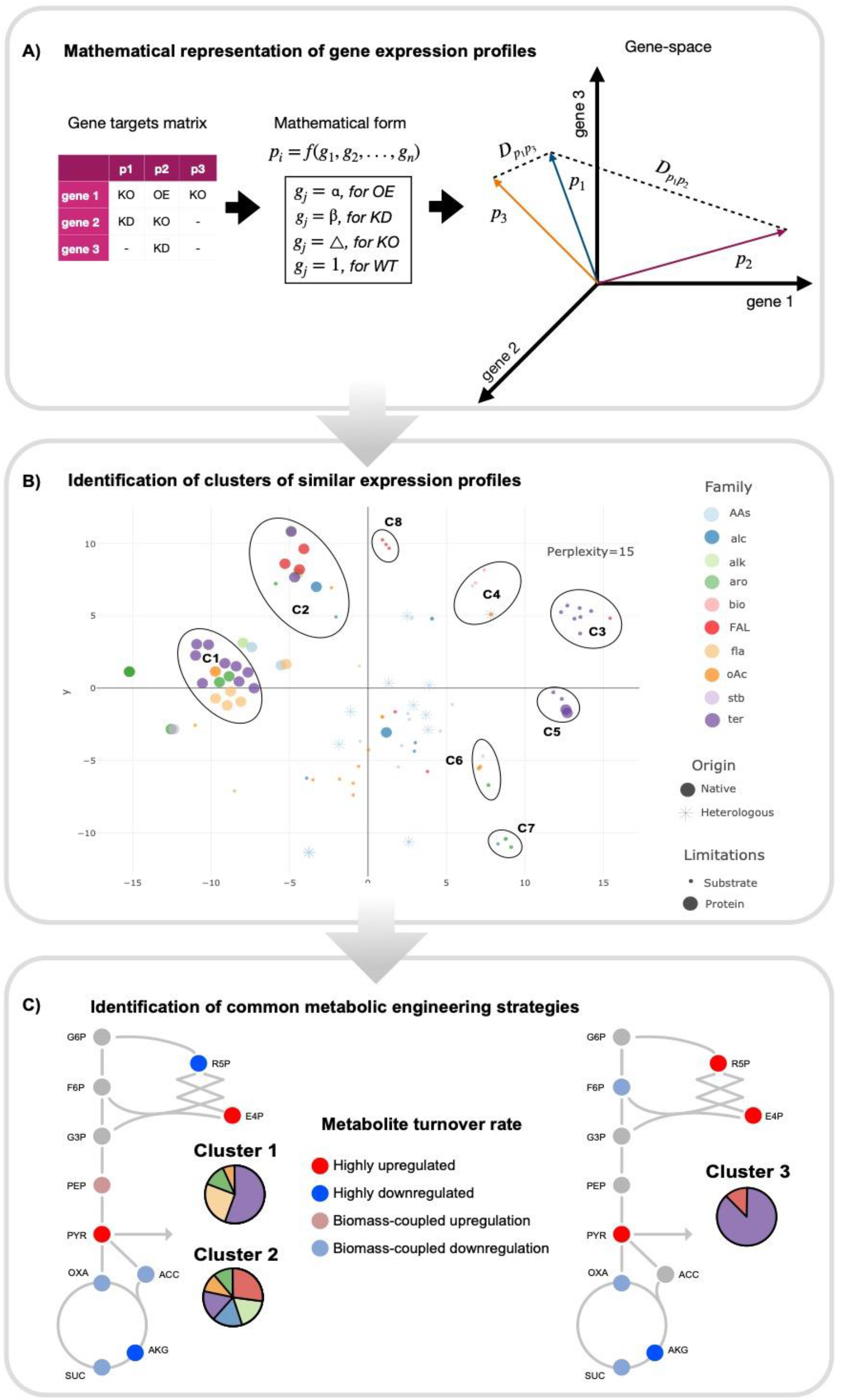
Model-driven design of platform strains for diverse chemical production. A) representation of gene targets for optimal production as mathematical vectors. B) Identification of clusters of products with similarities in their predicted engineering targets using t-SNE. Chemical families are indicated as AA, amino acids; Alc, alcohols; Alk, alkaloids; Aro, aromatics; Bio, bioamines; FAL, fatty acids and lipids; fla, flavonoids; oAc, organic acids; stb, stillbenoids; ter, terpenes. C) FBA predicts cluster-specific metabolic rewiring strategies. Fold-change in turnover rate of the main metabolic precursors, compared to wild-type, necessary for optimal production of the products in clusters 1, 2 and 3.

**Table 2.** Shared gene targets within each cluster of products.

| Cluster | Chemical Products | Shared KO targets | Shared KD targets | Shared OE targets |
| --- | --- | --- | --- | --- |
| 1 | betaxanthin, caffeic acid, vanillin β-glucoside, β-ionone, glycyrrhetic acid, miltiradiene, lycopene, taxadien-α-yl acetate, protopanaxadiol, genistein, quercetin, catechin, kaempferol, patchoulol, oleanolate, lupeol | RNR1, RNR4, RNR3, CAR2, FAA4, FAA1, FDH1 | SAH1, ARG5,6, MET6, LPD1, ADO1, MAE1, ARG7, MDH2, ARG8, ATP19 | NA |
| 2 | β-carotene, cinnamoyltropine, ARA, DHA, EPA, astaxanthin, psilocybin, docosanol | RNR1, RNR4, RNR3, CAR2, FAA4, FAA1, FDH1 | IDP1, ARG5,6, LPD1, MAE1, MDH1, ARG7, PPA2, MDH2, ARG8, ATP19 | NA |
| 3 | ergosterol, squalene, santalene, farnesene, amorphadiene, limonene, geraniol, artemisinic acid | NA | LPP1 | PDB1, PDA1, PDX1, ERG12, ERG8, LAT1, MVD1 |
| 4 | Itaconic acid, glutamine, proline, putrescine, spermine | NA | LPP1 | PDB1, PDA1, PDX1, LAT1 |
| 5 | valencene, nootkatone, linalool, β-amyryn | NA | ARG5,6, ARG8 | ERG12, ERG8, MVD1 |
| 6 | tryptophan, adipic acid, cis-muconate, hydroxymandelic acid | MAE1 | LPP1 | ARO4 |
| 7 | phenylalanine, 2-phenylethanol, mandelic acid, cinnamate | MAE1 | LPP1 | ARO4, ARO1, ARO2, SOL3, GND1, ZWF1, PHA2, ARO7 |
| 8 | Free-fatty acids, oleate, palmitoleate | NA | LPP1, ARG5,6, MAE1, CAR2, ARG8 | CDC19, BPL1, SOL3, GND1, PDC1, ACS2, PPA2, ZWF1, ACC1, ALD6 |

In general, these clusters are composed by products that belong to different chemical families, with the exception of cluster 3 and 5, composed mostly by terpenes, and cluster 8, formed just by lipid compounds. Mapping product origin (native or heterologous) and protein limitations information into the clustering results showed that, clusters 1 and 2 are composed by heterologous and highly protein-constrained products belonging to different compound classes; terpenes whose production is constrained by substrate availability tend to group together, in cluster 3; and most native products, despite their protein limitations, do not fall into the identified clusters. Altogether, this shows that metabolic engineering strategies for the different product clusters are defined by gene modifications that are related with redirecting flux and energy from central metabolism to the final specific heterologous pathways. This suggests that shared molecular characteristics between products (i.e., chemical classification of products) might not be the most decisive aspect when designing genetic modification strategies for enhanced production of multiple chemicals (platform or chassis strains).

In order to understand the particular metabolic rewiring required by each platform strain designed with the aid of the cluster analysis, turnover rates were calculated for the 12 main precursor metabolites in central carbon metabolism (D-glucose-6-phopshate, D-fructose-6-phosphate, ribose-5-phosphate, erythrose-4-phosphate, glyceraldehyde-3-phosphate, 3-phosphoglycerate, phosphoenolpyruvate, pyruvate, acetyl-CoA, 2-oxoglutarate, succinyl-CoA and oxaloacetate)^11^ using FBA simulations for different scenarios, optimal biomass formation and optimal production of each of the studied chemicals. Fold-changes were then computed for each of the precursor turnover rates, by comparing the optimal production flux distributions to their optimal biomass formation counterpart, for all 102 production scenarios. In this way fold-changes higher than one indicate that, for increased production, the overall flux towards a precursor should be upregulated, in comparison to a wild-type metabolic state, while fold-changes lower than one imply that the flux towards a precursor needs to be down-regulated (see **Materials and Methods** section in the **Supplementary Materials**).

**Figure 4C** shows that significant upregulation of flux towards erythrose-4-phosphate (E4P) and pyruvate, moderate upregulation of phosphoenolpyruvate (PEP), a drastic decrease in ribose-5-phosphate (R5P) and α-ketoglutarate (AKG) turnover rates and, a moderate down-regulation of the flux towards oxaloacetate (OXO), acetyl-CoA and succinyl-CoA should be combined to achieve optimal production levels of the products in clusters 1 and 2. Additionally, it can be seen that fluxes towards precursors located downstream from pyruvate (TCA cycle intermediates and acetyl-CoA) are needed to be downregulated for products in these clusters. This can be explained by a lower demand of building blocks, due to the decrease of biomass formation rate in a production scenario. Moreover, as all products in these clusters were found to be protein-limited, a predicted coordinated down-regulation of the lower section of central carbon metabolism suggests that forcing a fermentative regime, in which most of the energy is produced by glycolysis to minimize the protein burden induced by cellular respiration, thus, leaving room for expression of inefficient heterologous enzymes, offers the optimal conditions (metabolic mode) for production of these chemicals.

For the case of products in cluster 3, predictions indicate that a metabolic rewiring that induces significant upregulation of R5P, E4P and pyruvate production, and intense down-regulation of the flux towards and α-ketoglutarate is needed to improve production of these terpene compounds (**Figure 4D**), suggesting that an increased supply of NADPH (produced in the first steps of the pentose phosphate pathway, preceding ribose-5-phosphate) is needed for these products. The gene target profiles for the bioamines putrescine and spermine were found to cluster together with their precursor amino acids proline and glutamate, as well as itaconic acid (cluster 4). **Figure S7A** shows that genetic modifications common to all products in this cluster cause only moderate changes in the turnover rate of central carbon metabolism precursors, mostly for those in lower glycolysis, indicating that the optimal production mode for these products does not differ significantly from a wild-type optimal growing metabolic strategy. A strong requirement for increased flux towards E4P was found to be common to all terpenes, despite the protein limitations involved in their production pathways, as shown by Figures **4C**, **4D** and **S7B**.

Production of native and heterologous products derived from the shikimate pathway, those in clusters 6 and 7, were found to require an increase of flux towards the immediate precursors E4P and PEP, together with enhanced NADPH supply, provided by an increased flux to R5P, and a reduction of the metabolic turnover of precursors located downstream of PEP, in order to maximize carbon conversion (**Figure S7C**). Finally, significant increase of acetyl-CoA turnover, together with a moderate upregulation of the pentose-phosphate pathway for increased NADPH flux, was found to be the optimal reprogramming strategy for production of free fatty acids, oleate and palmitoleate (**Figure S7D**), resembling previous successful work in yeast cells^56^.

The set of common target predictions for a given cluster of products provides a modulated gene expression program capable of rewiring central carbon metabolism for increased production of key precursor metabolites. Implementation of these predictions in yeast cells can be used to drive the development of platform strains, specialized in providing the production scaffold for multiple chemicals. Platform strains can then be transformed into product-specific ones by introducing the necessary heterologous genetic components. This platform-based procedure will potentially reduce the resources and efforts involved in the development of next-generation cell factories.

### Web-based resources for exploration of metabolic engineering targets in S. cerevisiae

Predicted gene targets for increased production of the chemicals in this study were incorporated into metabolic atlas for visualization in a metabolic network context. **Figure 5** shows the gene modifications for improved patchoulol production in the central carbon metabolism of yeast as an example, where genes indicated for OE, KD and KO can be found. Furthermore, metabolic maps for other pathways, even in secondary and intermediate metabolism, are also available. Visualization options for the 102 products can be found at: www.dev.metabolicatlass.org. Additionally, in order to facilitate the utilization of the ecFactory method, interactive tutorials for prediction of engineering targets for 2-phenylethanol and heme production in yeast are available as MATLAB live scripts at: https://github.com/SysBioChalmers/ecFactory/tree/main/tutorials.

**Figure 5.**
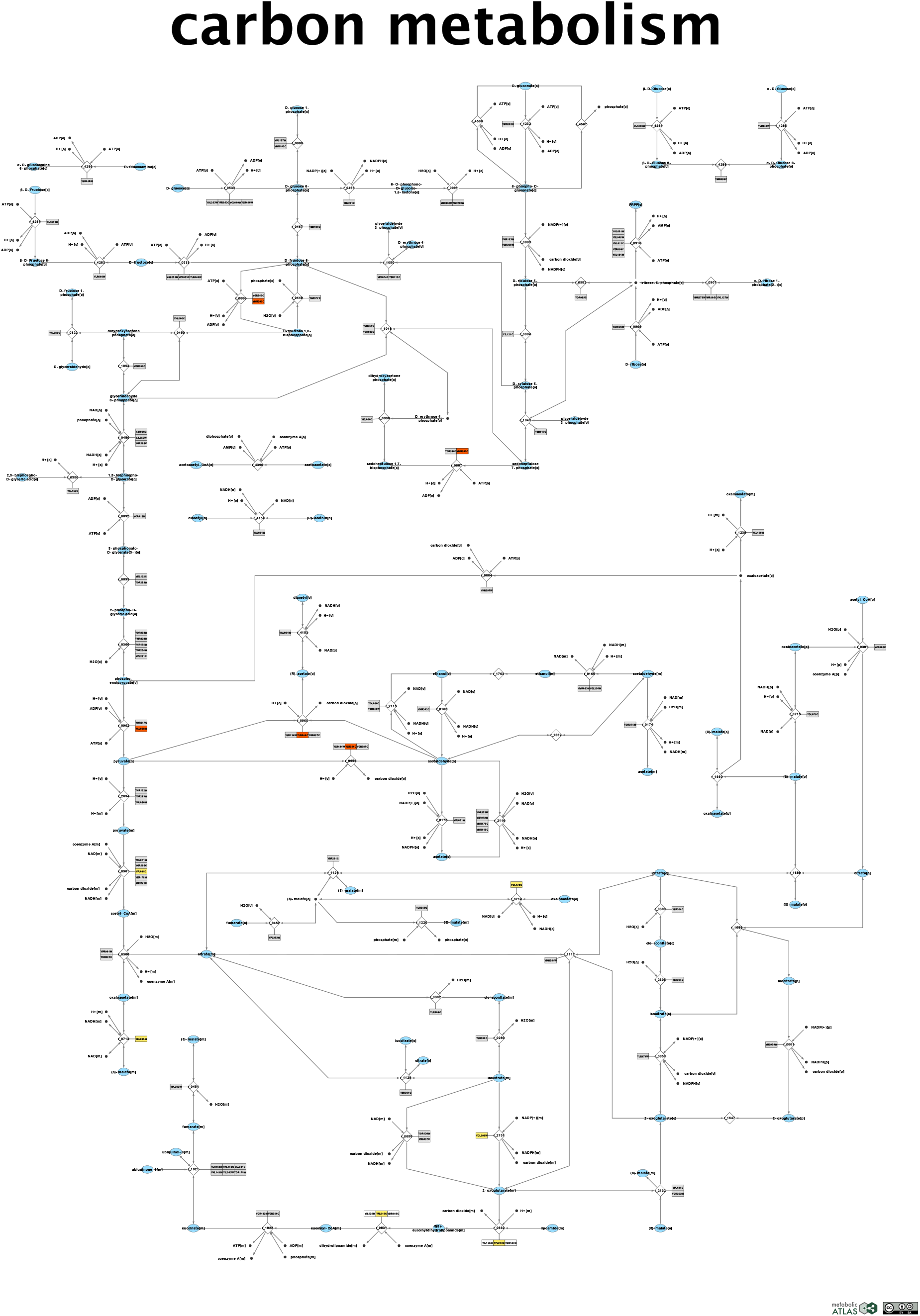
Map of *S. cerevisiae’s* central carbon metabolism from metabolic atlas. Gene targets for increased production of the terpene patchoulol are shown in red, for overexpressions; yellow for down-regulated targets; and gene for predicted gene deletions.

## Conclusions

Here we demonstrated that, by accounting for enzyme limitations, the use of metabolic models for quantitative prediction in metabolic engineering can be extended and improved. Enzyme-constrained models enabled assessment of the impact of enzyme capacity on the total protein and substrate costs of chemical production in cell factories, and reduction of the number of gene engineering targets for increased production predicted by stoichiometric constraint-based methods to a minimal optimal set of modifications. The model ecYeastGEM was used to predict gene engineering targets for enhanced production of 102 chemical products with yeast cells, including native and heterologous biochemicals with distinct chemical characteristics. Predictions showed to resemble complex engineering strategies that involve coordinated modulation and coordination of multiple pathways. Notably, supportive experimental evidence was found in the literature to verify the gene target predictions in 22 of the studied chemicals.

Sets of gene targets common across products were identified for 8 different groups of chemicals, inferred with a clustering algorithm. Flux balance analysis simulations indicate that, these core genetic modifications represent the expression tunning profiles, needed to rewire the central carbon metabolism of yeast towards increased production of the main metabolic precursors required by groups of valuable chemicals. By visualizing the 8 different rewiring schemes we learned that clustering of products according to their gene target predictions obeys to combinations of these three basic factors: 1) protein burden induced by the specific production pathways and its impact on energy production; 2) the metabolic precursor that provides the main carbon flux for final product formation; 3) products that require increased NADPH flux levels. Thus, the presented approach suggests the advantages of using of enzyme-constrained models for design and understanding of platform strains optimized for diverse chemical production. Nonetheless, expanding the scope and number of chemicals and host organisms for this kind of large-scale studies might help to unveil additional core principles for rationally engineering of metabolism.

We envision that the tools and methodology developed in this study will contribute to accelerate development of robust and efficient microbial strains both for specialized and also versatile production of valuable chemicals, promoting the conversion from petrol a bio-based economy.

## Acknowledgments

This project has received funding from the Novo Nordisk Foundation (grant no. NNF10CC1016517), the Knut and Alice Wallenberg Foundation, and the European Union’s Horizon 2020 research and innovation program with projects DD-DeCaF and CHASSY (grant agreements No 686070 and 720824). This project was also supported by The Shanghai Pujiang Program and Grants 22208211 from the National Natural Science Foundation of China (NSFC).

## Author Contributions

I.D. wrote the draft manuscript, developed the software and methods and analyzed results. Y.L. performed the literature review, constructed the heterologous pathways in ecYeastGEM and contributed to method development. All authors reviewed and edited the manuscript. H.L. and J.N. designed and conceived the initial project. J. N. and J. S. obtained the funding for this study.

