## Supplementary Material for "Computational biology predicts metabolic engineering targets for increased production of 102 valuable chemicals in yeast"

### Supplementary materials

Table of content

Materials and Methods …………………………2
Supplementary figures …… …………………..14
Supplementary tables …………………………. 23

### 1.- Materials and Methods

#### 1.1 Incorporation of heterologous metabolic pathways into ecYeastGEM

The metabolic pathways for synthesis of heterologous products in *S. cerevisiae* cells were incorporated into ecYeastGEM based on engineered pathways reported by independent studies available in the literature. For those cases with multiple developed pathways, those with highest yield or production rate were selected (see **supp. table S2** for a complete list of references to the selected study cases). The incorporation process mainly included adding new reactions, updating existing reactions, genetic manipulation, and setting constraints. New reactions of each product were separately added into ecYeastGEM following the GECKO approach[1]. All $k_{cat}$ values were manually collected from the BRENDA database[2] and scientific literature, together with the corresponding enzymatic molecular weights (kDa).

Additionally, missing or modified genes, proteins and $k_{cat}$ values were added to preexisting reactions, when needed, and named as 'updated' together with the original RxnID in the model. In order to simulate alleviation of feedback inhibition by mutation at specific site for some especific genes, such as *Aro3*, *Aro4*, and *Aro7*, the original enzymes were substituted by heterologous ones ($Aro4_{K229L}$, $Aro4_{G226S}$, $Aro7_{T226I}$, and $Aro7_{G141S}$) with higher $k_{cat}$ value.

#### 1.2 Computing production capabilities using GEMs and ecModels

The feasible production space for each of the chemicals of interest for this study were explored by quantifying maximum production yields at varying levels of biomass yield, spanning from its maximum value (0.48 $g_{biomass}/g_{glucose}$, according to both YeastGEM and ecYeastGEM[3]) to a value of zero, a situation in which all the carbon is used for product formation. For both model types and all chemicals a series of 10 flux balance analysis problems (FBA) were solved[4], using a fixed unit glucose uptake rate, unlimited uptake of other nutrients available in minimal mineral media YPD, and fixed varying levels of suboptimal biomass production. This series of FBA problems are expressed in the following mathematical form:

$For\ i = 1, \dots., n$

$$max. \quad C^T v = v_p$$

$subject\ to$

$$Sv = 0$$

$$LB < v < UB$$

$$v_{glc}^{in} = v_{glc}^{Ex}$$

$$v_{biomass} = \left(\frac{i-1}{n}\right) v_{biomass}^{MAX_1}$$

Where $v$ represents the vector of reaction rate fluxes in the metabolic model; $S$, the stoichiometric matrix containing the stoichiometric coefficients of each metabolite (rows) in their corresponding reactions (columns), negative values indicate consumption, whilst positive values indicate production of a metabolite; $C^T$ is the transposed vector of coefficients for each flux variable in the linear objective function; $v_p$ represents the production flux for the product of interest ($p$); $LB$ and $UB$ represent the vectors of lower and upper bounds for each of the metabolic fluxes, respectively; $v_{glc}^{in}$ is the glucose uptake reaction; $v_{glc}^{Ex}$, is the fixed value used for each case, 1 mmol/gDw h and 10 mmol/gDw h, for low and high glucose regimes, respectively; $v_{biomass}$ is the biomass production pseudo-reaction; and $v_{biomass}^{MAX_1}$ is the maximum biomass production rate attainable with a unit glucose uptake rate.

Biomass formation and production rate were both divided by the fixed glucose uptake rate in order to transform them to yield quantities. Both yield vectors are mapped in a two-dimensional space to generate yield plots, in which the unfeasible production regions, according to the enzyme capacities profile in the cell, can be easily identified. Yield plots for multiple products are shown in **Fig. 1D-E** and **Fig. S1**.

#### 1.3 Computing substrate and protein cost of biochemical production using ecModels

Optimal flux distributions were obtained for all the 102 products by running FBA simulations with ecYeastGEM. The respective product secretion reaction was used as objective function for each case, unconstrained uptake of mineral minimal media components was allowed and glucose uptake rate was constrained with an upper bound of 1 mmol/gDw h.

From these flux distributions minimal substrate costs were calculated as:

$$C_{S_i} = \frac{v_{glc}^{in} * MW_{glc}}{v_i^{opt} * MW_i}$$

Where $v_{glc}^{in}$, represents the effective glucose uptake rate (lower than 1 mmol/gDw h for highly protein-constrained products); $MW_{glc}$ is the molecular weight of glucose (0.18 g/mmol); $v_i^{opt}$ represents the optimal production rate for *i-th* metabolite ; and $MW_i$, its molecular weight.

Minimal protein costs were calculated as:

$$C_{P_i} = \frac{\sum_j MW_j e_j}{v_i^{opt} * MW_i}$$

Where $\sum_j MW_j e_j$, represents the sum of the product of each enzyme usage reaction ($e_j$, in mmol/gDw) by its molecular weight ($MW_j$, in g/mmol of enzyme); $v_i^{opt}$ represents the optimal production rate for *i-th* metabolite ; and $MW_i$, its molecular weight.

#### 1.4 ecFactory: a multi-step method for prediction of metabolic engineering gene targets

The ecFactory method is a series of sequential steps for identification of metabolic engineering gene targets. These targets show which genes should be subject to overexpression, modulated expression (knock-down) or deletion (knock-out), with the objective of increasing production of a given metabolite. This method was developed by combining the principles of the FSEOF algorithm (flux scanning with enforced objective function)[5] together with the features of GECKO enzyme-constrained metabolic models (ecModels)[1], which incorporate enzymes as part of genome-scale metabolic networks. All the sequential steps of this method are included in the main function ***run_ecFactory*** and are explained in the following numbered sections.

#### *1.4.1 FSEOF for ecModels*

As a first step, the ecFactory pipeline runs the function ***run_ecFSEOF***, available in the utilities folder of the GECKO toolbox[6]. This is an implementation of the FSEOF algorithm for a specified cellular objective (e.g. secretion rate of a given metabolite) suited for the characteristics and format of ecModels. The algorithm solves a series of 16 different FBA problems, using fixed biomass yield as constraints at regularly decreasing intervals, spanning from 100% to 25% of the maximum predicted value (0.48 $g_{biomass}/g_{glucose}$ for ecYeastGEM v8.3.4). Reactions whose flux increases or decreases consistently across the resulting 16 optimal flux distributions are identified as reaction step targets for engineering. For each of these reactions, an overall flux score is calculated as:

$$V_{score_i} = \frac{\sum_{j=1}^{n} v_{ij}}{n\left(v_i^{bio_{MAX}}\right)}$$

Where $v_i^{bio_{MAX}}$ represents the flux carried by reaction *i* at the maximum biomass yield condition; and $\frac{\sum_{j=1}^{n} v_{ij}}{n}$, the average flux carried by reaction *i* across all flux distributions (*n*=16 steps). In contrast to classic *FSEOF*, flux scores are computed as rescaled mean flux values across the spanned conditions, instead of considering a constant slope in between the initial and final simulation scenarios, due to possible changes of metabolic regime (respiratory to mixed fermentative) while spanning suboptimal biomass yields. Any $V_{score_i}$ value higher than 1000 is truncated to 1000, in order to avoid infinity values. Undetermined values (zero divided by zero) are assigned with a value of 1. Then, gene scores, referred to as *K-scores* from here onwards, are computed as the average of reaction flux scores ($V_{score_i}$) across all the reactions that are encoded by each gene:

$$K_{score_i} = \frac{\sum_{j=1}^{m} V_{score_j}}{m}$$

Where $K_{score_i}$ is the score for gene target *i*; $\sum_{j=1}^{m} V_{score_j}$ is the sum of the reaction scores of all reactions that are catalysed by gene product *i*; *m* is the total number of reactions catalysed by gene product *i*. Genes with a *K-score* higher than 1 are suggested as targets for overexpression; genes with 0.05 < *K-scores* < 0.5 as targets for knock-down; and those with *K-scores* < 0.05 as knock-out targets.

#### *1.4.2. Identification of flux leaks*

In the FSEOF algorithm, gene engineering targets are identified based on the assumption that there is a trade-off between biomass formation and product secretion or accumulation. Nevertheless, additional ways of enhancing production can be found by focusing on other characteristics of the metabolic network. Reaction steps that consume the product of interest, which may decrease accumulation or secretion, are often elusive to the FSEOF algorithm. These reactions might not be relevant for biomass formation, especially for those located in metabolic pathways outside of central carbon metabolism. In order to capture this kind of reactions in the list of gene target candidates, all genes that encode for enzymes catalyzing these reaction steps, possibly being detrimental, are added to the list of  candidates by the funciont ***find_flux_leaks***.

#### *1.4.3. Discarding essential genes from deletion targets*

Experimentally validated essential genes (retrieved from the *Saccharomyces* genome deletion project, available at: http://www-sequence.stanford.edu/group/yeast_deletion_project/downloads.html) are discarded from the list of knock-down and knock targets.

#### *1.4.4. Identification of enzyme groups in the candidate targets*

As ecModels are capable of capturing complex reaction-gene relations, isoenzymes and enzyme complexes, some of the gene products predicted as targets by FSEOF may fall into one of these categories. Isoenzymes and enzyme complex subunits are identified by finding all groups of identical vectors in a metabolite-gene matrix, which connects all genes with the metabolites involved in the reactions catalysed by their corresponding enzymes. In an ecModel, the metabolite-gene matrix is defined as:

$$MG = logical(S) * RxnGeneMat$$

Where $logical(S)$ is the Boolean form of the model's stoichiometric matrix, representing all metabolites as rows and reactions as columns; $RxnGeneMat$, is a Boolean matrix that establishes the relationship between enzyme-encoding genes and biochemical reactions in the ecModel, in which a value of 1 indicates genes whose protein products catalyze a given reaction. The $MG$ matrix is computed by the function ***getMetGeneMatrix*** which feeds it into ***getGeneGroups***, a function that finds groups of enzymes related to exactly the same metabolites and also identifies all enzymes that are unique, in their metabolites association, from the list of targets. This information is used later in the ecFactory pipeline in order to decide which genes should be modified simultaneously for a given objective.

#### *1.4.5. Computing enzyme demands with flux variability analysis*

In order to compute enzymatic demands for each of the gene candidates under different production scenarios, flux variability analysis (FVA) is run for the corresponding enzyme usage reactions in the ecModel. A feasible enzyme usage variability range is defined as:

$$\Delta_{e_i} = e_i^{max} - e_i^{min}$$

The values for $e_i^{max}$ and $e_i^{min}$, are obtained by optimizing (maximization and minimization) the following linear programming problem:

$$\text{Min or Max. } z = e_i \quad \forall \ i = 1, \dots, m$$
$$\text{s.t. } Sv = 0,$$
$$LB_j \leq v_j \leq UB_j, \qquad j = 1, \dots, n$$

Where $e_i$ represents the usage reaction for enzyme $i$; $m$ is the number of metabolic enzymes associated to the gene target candidates; $S$ is the stoichiometric matrix of the metabolic network, representing metabolites in its rows and reactions in its columns, negative coefficients are assigned to reaction substrates whilst positive signs are assigned to products; $v$, is the vector of reaction fluxes in the network; $LB_j$ is the lower bound value for reaction $j$; $UB_j$ is the lower bound value for reaction $j$ and $n$ is the total number of reactions in the model. In the ecFactory pipeline, enzyme usage variability range profiles are obtained for two different scenarios:

1) Fixed glucose uptake rate of 1 mmol/gDw h, fixed biomass production rate, equal to the maximum attainable value under this glucose uptake ($v_{bio}^{max}$). Which represents the enzyme demand ranges for an optimal growing strain (growth scenario).

2) Fixed glucose uptake rate of 1 mmol/gDw h, a lower bound of 0.5* $v_{bio}^{max}$ for the biomass production pseudoreaction and a fixed value of $v_{prod}^{max}$ for the production reaction of interest, which corresponds to the maximum attainable production value under a unit glucose uptake rate and the specified biomass production. The variability ranges obtained by this represent profiles of enzyme demands for the predicted targets in an enhanced production scenario.

This analysis is run by the function ***enzymeUsage_FVA***, part of the ***ecFactory*** pipeline. Additionally, a parsimonious flux distribution is obtained by minimizing the total enzyme mass demand (protein pool usage pseudo-reaction), while subjected to the same set of constraints used for each of the conditions specified above.

***1.4.6. Reducing the list of candidates according to enzyme variability ranges***

Enzyme usage variability ranges for the two aforementioned scenarios are used to classify the enzyme targets. **Figure S4** illustrates a reduced network, with the metabolic task of reaching node D starting from A, in which the second enzymatic step (B $\rightarrow$ C) can be catalised by three different isoenzymes ($E_{2A}$, $E_{2B}$ and $E_{2C}$). In this case, any of the three isoforms can be used for production of D, therefore the three enzymes display a usage variability range that spans from zero to a specific maximum value (occurring when all the reaction flux is carried by a single isoform). Nonetheless, in parsimonious flux distributions (those that minimize the total sum of fluxes in the solution) usually just the most efficient isoenzyme of a given group will carry a non-zero flux[7], in this case very close to its maximum feasible value, therefore, in ecFactory such enzymes are classified as "optimal for production", whilst the other isoforms as "suboptimal for production". $E_3$ represents a typical case of an "essential" enzyme, whose usage flux cannot take a zero value and its whole range is confined to a region close to its maximum feasible value. The network shows that $E_4$ is a "futile" enzyme for achieving the metabolic task of producing D, therefore its maximum, minimum and parsimonious flux are all equal to zero.

This classification of enzymes can be based on the profiles of variability ranges both for optimal production and optimal growth scenarios, additionally, provides the following criteria for reduction of the number of gene targets predicted by the ecFactory method:

- Discard OE target candidates classified as futile for production
- Discard OE target candidates classified as suboptimal for production, focusing on optimal isoforms when needed.
- Discard enzymes essential for production from KO target candidates.
- Discard isoenzyme groups that contain an optimal isoform for biomass formation from KD and KO target candidates. This aims to reduce mutant complexity, as in order to have an effect in redirecting flux from biomass to production, coordinated modification of all the isoenzymes in the group would be necessary. Focusing on essential enzymes for biomass production is recommended for KD targets and futile ones for production for KO targets.

For each enzyme in the list of candidate targets, comparison in between the two different demand scenarios enables classification of differential demand patterns as: distinct, overlapped and undistinguishable (Figure S5). Enzymes with undistinguishable demand patterns (Figure S5C and S5F) are discarded from the candidates list. Moreover, enzymes with distinct demand patterns are assigned a degree of priority of 1 whilst a value of 2 is assigned to those with overlapped demand patterns, in order to inform experimental design for testing modifications.

#### *1.4.7. Finding an optimal combination of genetic modifications*

The enzyme usage ranges under the production scenario for the candidate targets provide an approximation to enzyme expression levels characteristic of an optimal producing strain. In order to simulate this optimal behaviour, each of the enzymes in the list of remaining targets are constrained according to the following:

$$e_{i_p}^{Prod} \leq e_i \leq e_{i_{max}}^{Prod}, \qquad i = 1, \dots, n$$

Where $e_i$ is the usage reaction for enzyme *i*; $e_{i_p}^{Prod}$ is the parsimonious usage value for enzyme *i*, subjected to the optimal production scenario constraints; $e_{i_{max}}^{Prod}$ is the maximum feasible usage value for enzyme *i*, subjected to the optimal production scenario constraints; and $n$ represents the number of gene targets with an associated enzyme in the ecModel. The suboptimal biomass production value, used as lower bound for the enzyme usage variability analysis in the optimal production scenario, is also used as a constraint in this step, then the optimal production rate is computed by maximization of its corresponding reaction rate. Once a maximum production value has been obtained, it is used to fix both the lower and upper bounds for its corresponding reaction and the glucose uptake rate is then minimized, in order to compute a maximum optimal product yield. These production values are representative of an optimal strain, in which expression levels of key enzymes (encoded by the gene candidate targets) have been fine-tunned (OE, KD or KO), and differentiated from those in the optimal biomass formation scenario (wild-type), in order to redirect flux towards the product of interest. However, as metabolism is highly interconnected and kinetic parameters of enzymes are heterogenous in nature, it is possible that not all of the constrained enzyme targets contribute to achieve the optimal production phenotype. To investigate this, the constraints for each of the enzyme targets are "returned" to their optimal biomass producing levels (wild-type expression) as follows:

$$e_{i_p}^{Bio} \leq e_i \leq e_{i_{max}}^{Bio}, \qquad \forall\, i = 1, \dots, n$$

Where $e_{i_p}^{Bio}$ represents the parsimonious usage of enzyme *i* in the optimal biomass formation scenario; $e_{i_{max}}^{Bio}$ is the maximum feasible value for the usage of enzyme *i* under the same conditions. Enzymes are modified independently in an iterative procedure, computing the following evaluation metric at each step:

$$\eta_i = \frac{1}{2}\frac{v_P^{mod}}{v_P^{Opt}}\left(1 + \frac{v_{glc}^{Opt}}{v_{glc}^{mod}}\right)$$

Where $v_P^{mod}$ represents the maximum production value for the product of interest, obtained when $e_i$ has been "returned" to its wild-type demand values, while keeping the rest of enzyme targets on their optimal production ranges; $v_P^{Opt}$ is the maximum production rate for the optimal production scenario; $v_{glc}^{mod}$ is the minimum glucose uptake rate obtained when $e_i$ has been "returned" to its wild-type demand values, while keeping the rest on their optimal production ranges; $v_{glc}^{Opt}$ is the minimum glucose uptake rate under the optimal production scenario conditions. A $\eta_i$<1 indicates that modified expression levels of enzyme $i$, in comparison to the wild-type enzyme allocation profile, are needed to achieve the optimal production scenario, therefore the gene encoding for enzyme *i* is kept in the list of target candidates. Values $\eta_i$=1 indicate that modification of the expression levels of enzyme *i* is not necessary to reach optimal production, therefore the gene encoding for it would be removed from the list of candidate targets, and the bound for its respective enzyme usages are kept as those needed for optimal biomass formation (wild-type) for the rest of the iterations.

This iterative approach assesses the effects of combined genetic modifications in the simulated phenotype, by comparing performance under two different enzyme usage profiles instead of relying on imposing arbitrary values for overexpression or knock-down levels over the bounds of enzyme usage reactions. All these steps are performed by the function *constructMinimalMutant* in the ecFactory pipeline, which returns an optimal mutant with the minimal subset of genetic modifications to achieve the optimal production scenario and a list describing each of the successfully combined genetic modifications. Moreover, the subtractive approach of removing non-essential modifications from an optimal mutant ensures robustness of results, regarding the order in which each modification is tested. This is a major improvement in the method, in contrast to its previous version, with a cumulative approach to "optimal" strain construction, which is highly dependent on the order in which modifications are implemented into the model. The analysis of randomized test of modifications on the results can be explored in the following MATLAB live script available at:
https://github.com/SysBioChalmers/ecFactory/blob/feat/test_targets_randomization/tutorials/test_randperm.mlx

### 1.5 Computation of genetic target profile vectors

For each of the analysed chemicals, the list of all individually validated targets was mapped to a gene-space, gene target profile vectors are created by assigning a value of 4 for overexpression targets, 0.25 for knock-downs, 0 for deletions and a value of 1 for the rest of the genes present in the ecModel. These values were established according to the characteristic rescaling of the biomass-coupled reactions when switching from optimal biomass to product formation in the FSEOF part of the method. Column-wise concatenation of these target vectors returned a gene targets matrix ($\boldsymbol{G}$), therefore, this matrix offers a straightforward relation between genes (rows) and the studied chemical products (rows) for further analysis of results.

As the gene profile vectors live in an Euclidean space, pairwise distances can be computed between each pair of profile vectors, which gives a measure of how different are the genetic modification profiles for two different products. Product pairs with gene target vectors close to each other suggest the existence of gene targets in common, i.e. gene modifications that might enhance production of both products. A null distance between two different profile vectors indicates that are the same genetic modifications are predicted to enhance production of their respective chemical products. Gene targets pairwise distances were then calculated for all pairs of products (10,816 combinations) and organized in a matrix form ($\boldsymbol{D_G}$). All values in the $\boldsymbol{D_G}$ matrix where then normalized by the maximum value available in it, returning a matrix of continuous values in the [0,1] interval. A graphical representation of the mapping of gene target vectors and the pairwise distances between them is shown in **Fig. S6**.

#### 1.6 Computation of optimal flux vectors

Optimal production flux vectors were obtained for each of the studied chemicals by FBA simulations with the ecYeastGEM, using the respective production target as objective function, and a fixed unit glucose uptake rate with unconstrained uptake of minimal media components (YPD) as constraints. No biomass formation was forced in these simulations, in order to obtain flux vectors that reflect the path followed for optimal conversion of nutrients to final products. All the different production vectors are defined within the feasible solution space of ecYeastGEM, bounded by stoichiometric and enzymatic constraints.

#### 1.7 Cluster analysis of gene target profiles

Dimensionality reduction of the $\boldsymbol{G}$ matrix was performed using the t-SNE method in order to find clusters of similar vectors of genetic modifications across groups of chemicals. This was done by using the package R package *Rtsne*[8], using a fixed seed value for reproducibility of results. The perplexity hyperparameter was varied 2 to 25, which is a measure of the weight of local vs. global effects in the expected clusters[9]. Results of tSNE projected in a two-dimensional space, showed identifiable clusters of target vectors for specific products, that were consistent across the whole range of tested values of perplexity. Common gene targets (for OE, KD and KO) across all products in a given cluster were then found by analyzing the ***G*** matrix.

### 1.8 Metabolic precursors analysis

In order to assess the requirements for the 12 core metabolic precursors (glucose-6-phosphate, fructose-6-phosphate, ribose-5-phosphate, erythrose-4-phosphate, glyceraldehyde-3-phosphate, 3-phosphoglycerate, phosphoenolpyruvate, pyruvate, acetyl-CoA, 2-oxoglutarate, succinyl-CoA, oxaloacetate)[10] for each of the analyzed chemical products, metabolite turnover, or flux-sum, were computed for each of them according to the definition[11]:

$$\varphi_i = \frac{1}{2}\left|\sum_j^n S_{ij}^c v_j\right|$$

Where $v_j$ represents each of the reaction fluxes connected to metabolite $i$ (either production or consumption), $S_{ij}$ are the stoichiometric coefficients for metabolite $i$ in each of the corresponding reactions. This number represents the rate of transformation for internal metabolites at steady-state. Metabolic turnovers were calculated for the 12 precursors at a wild-type scenario (using the parsimonious flux distribution for optimal biomass production) and also for each of the 102 different optimal production scenarios described above. Computation of fold-change of metabolic turnover between optimal production and optimal biomass conditions were performed for the 102 chemical products in this study.

### 2.- Supplementary Figures

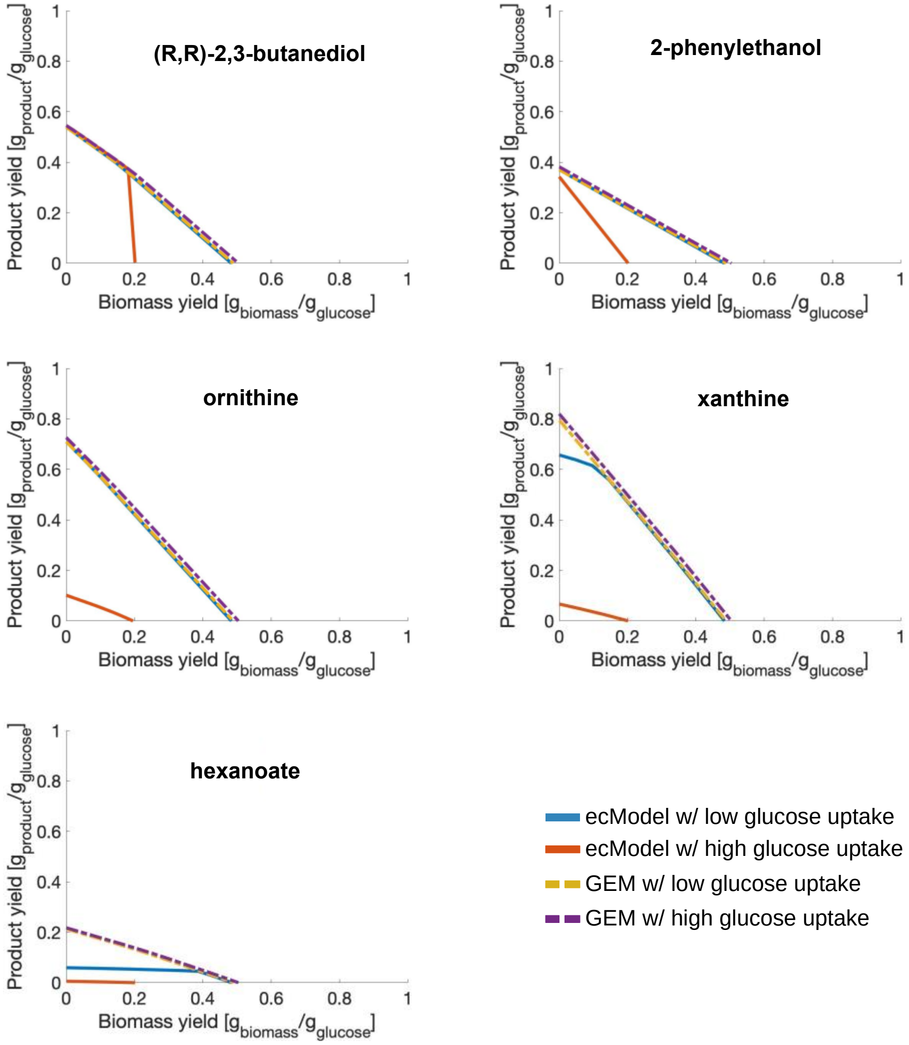


**Figure S1.-** Production yield space for selected native metabolites in yeast with different degree of enzymatic limitations.

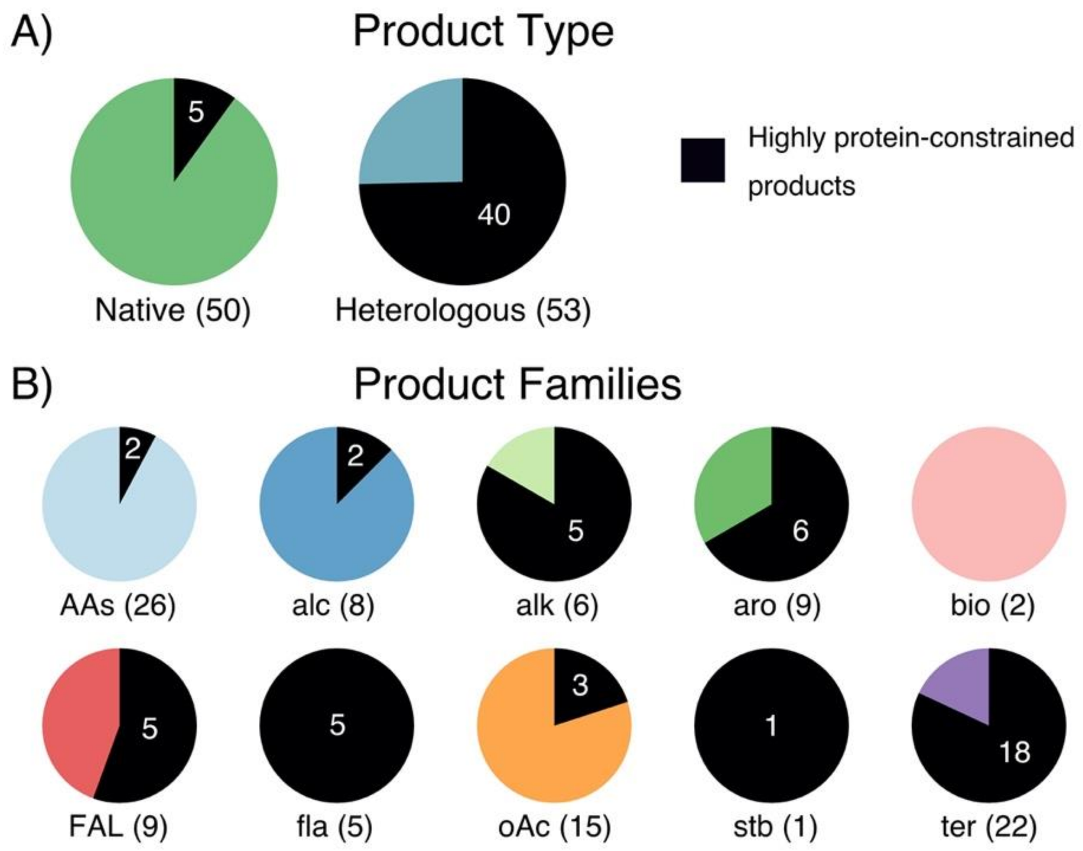


**Figure S2.-** Classification of 102 chemical products studied with ecYeastGEM. **A)** Distribution of chemicals divided by origin (native or heterologous), according to *S. cerevisiae's* metabolic network. **B)** Number of products per chemical family. Black sections and white numbers indicate the amount of products whose production was found to be protein-limited at low glucose uptake rates (1 mmol/gDw h).

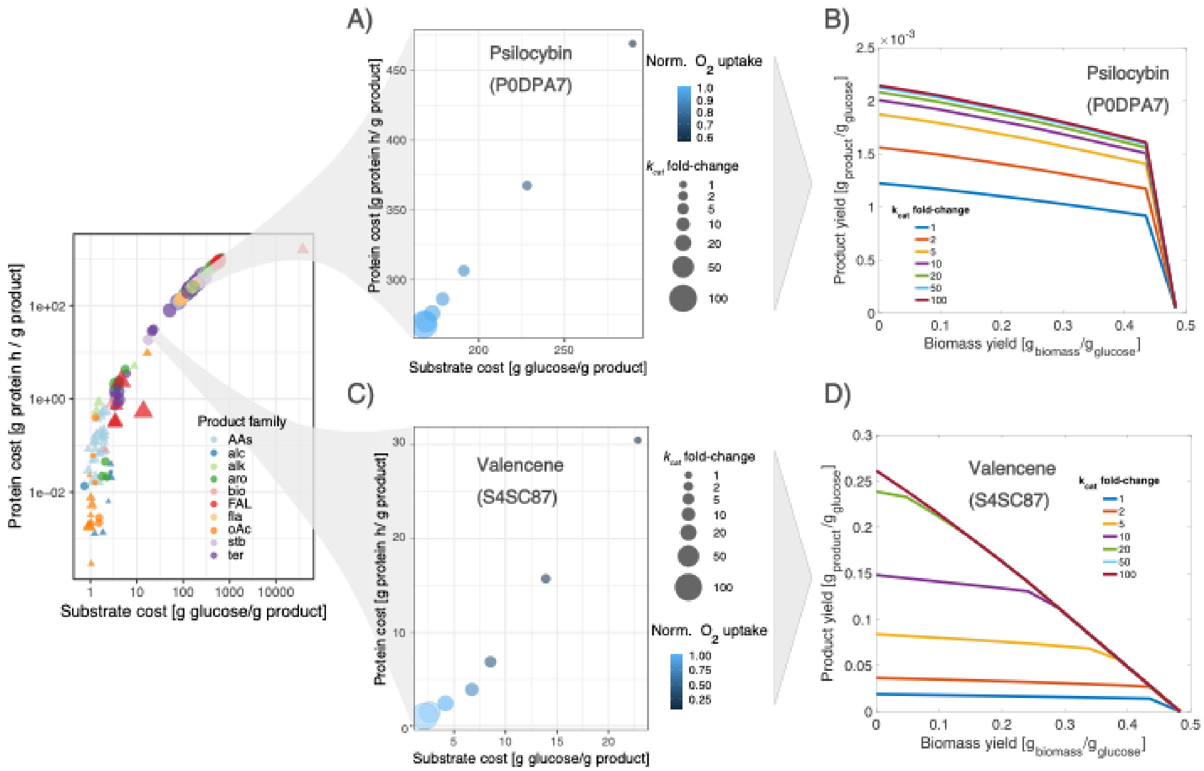


**Figure S3.-** Enzyme capacity and production capabilities. **A)** Effect of P0DPA7 (tryptamine 4-monooxygenase) enzyme capacity on the production cost for psilocybin production. **B)** Psilocybin production space predicted by ecYeastGEM. **C)** Effect of S4SC87 (terpene synthase) enzyme capacity on the valencene production space. **D)** Valencene production space predicted by ecYeastGEM.

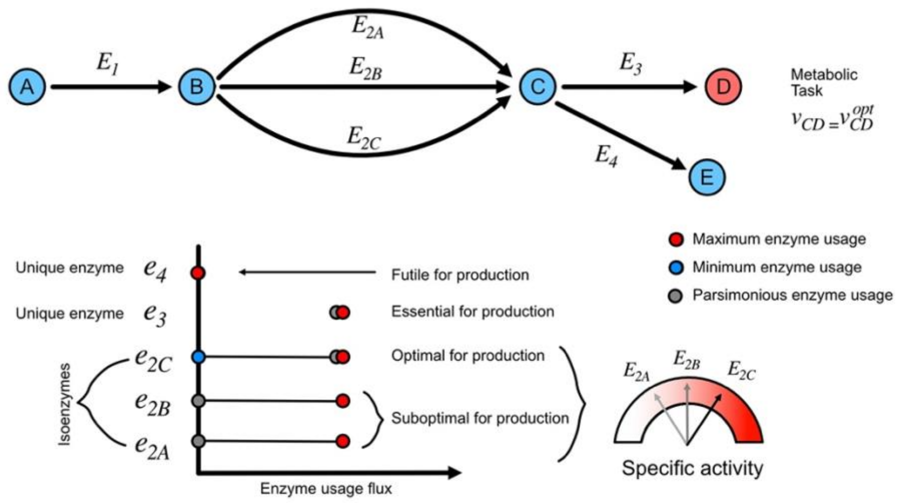


**Figure S4.-** Classification of gene targets according to their enzyme usage variability ranges. Upper section represents a simple metabolic pathway with enzymatic redundancy (presence of isoenzymes). Lower section represents a classification of enzymes ($E_i$)in the pathway according to the variability ranges of their respective enzyme usages ($e_i$) allowed by a fixed metabolic objective.

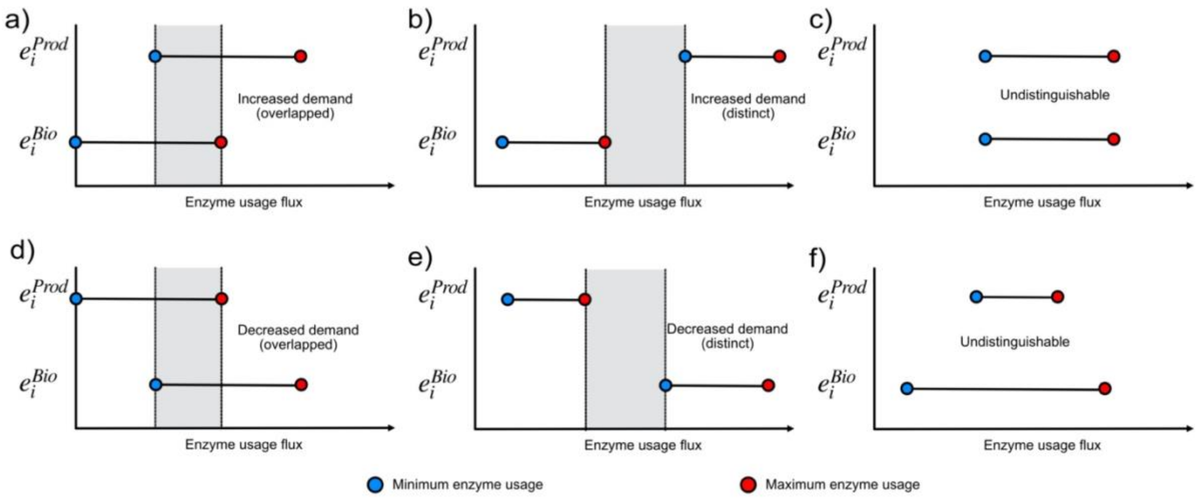


**Figure S5.-** Enzyme usage variability ranges under two scenarios: 1) maximization of product formation, while subject to a suboptimal biomass production, and 2) maximization of biomass formation subject to a minimal product formation. A) Increased enzyme demands with an overlapped region between objectives. B) Increased enzyme demands with distinct ranges in between objectives. C) Undistinguishable demand patterns between the two objectives, equal ranges. D) Decreased enzyme demands overlap with overlap between objectives. E) Decreased enzyme usage with distinct demand ranges in between objectives. F) Undistinguishable demand ranges, enzyme usage range for optimal production is a subset of the usage range for optimal biomass formation.

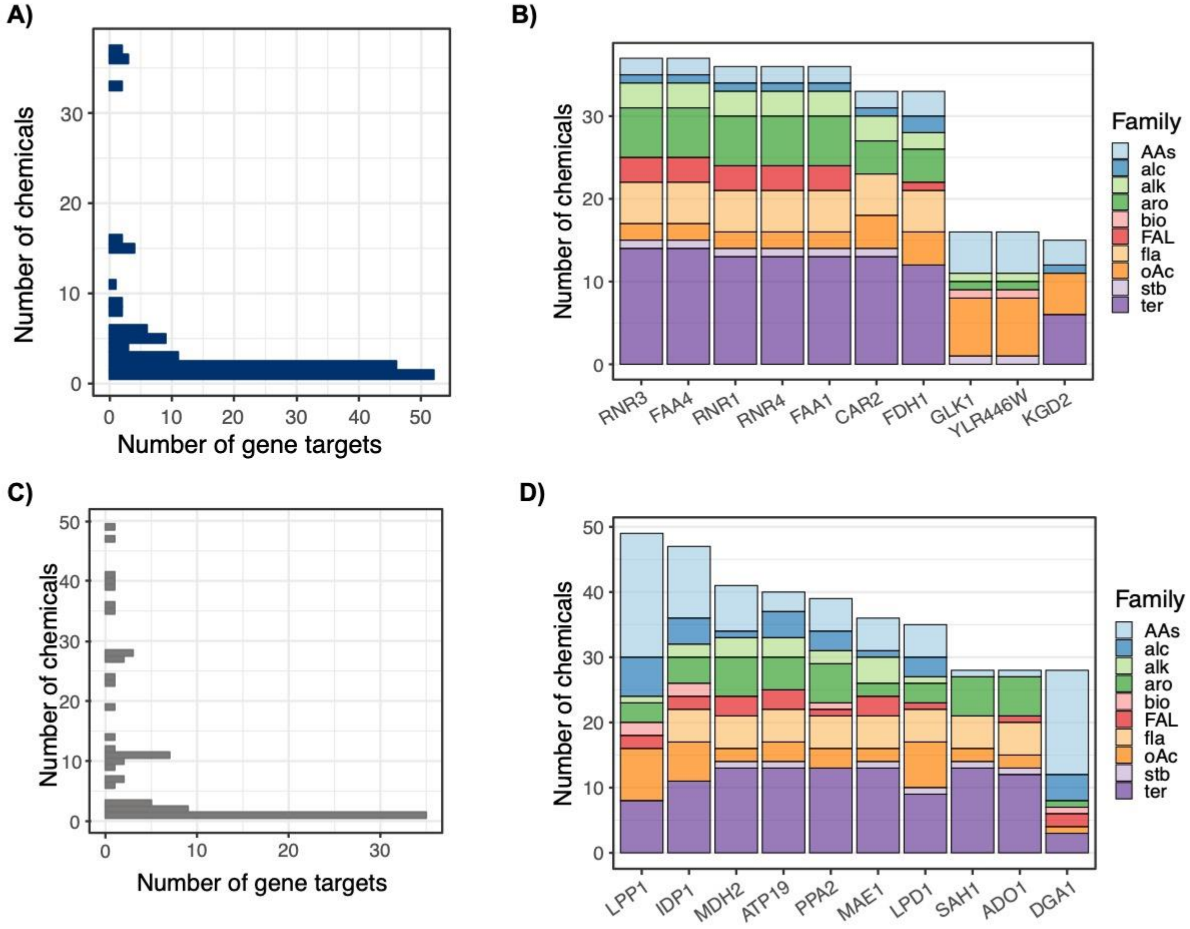


**Figure S6.-** Analysis of predictions for KO and KD gene targets across 102 chemicals. **A)** Distribution of the degree of product-specificity for KO gene targets across all chemicals. **B)** Top 10 most common gene KO tartets across products and chemical classes. Short gene names correspond to ribonucleoside-diphosphate reductase large chain 2 (RNR3), long-chain-fatty-acid—CoA ligase 4 (FAA4), ribonucleoside-diphosphate reductase large chain 1 (RNR1), ribonucleoside-diphosphate reductase small chain 2 (RNR4), long-chain-fatty-acid—CoA ligase 1 (FAA1), ornithine aminotransferase (CAR2), formate dehydrogenase 1 (FDH1), glucokinase 1 (GLK1), putative hexokinase (YLR446W), and multifunctional 2-oxoglutarate metabolism enzyme (KGD2). **C)** Distribution of the degree of product-specificity for KD gene targets across all chemicals. **D)** Top 10 most common gene KO targets across products and chemical classes. Short gene names indicate lipid phosphate phosphatase 1 (LPP1), isocitrate dehydrogenase [NADP] (IDP1), cytoplasmic malate dehydrogenase (MDH2), ATP synthase subunit K (ATP19), mitochondrial inorganic pyrophosphatase (PPA2), NAD-dependent malic enzyme (MAE1), mitochondrial dihydrolipoyl dehydronase (LPD1), adenosylhomocysteinase (SAH1), adenosine kinase (ADO1), diacylglycerol O-acyltransferase 1 (DGA1). Abbreviations for chemical family names indicate amino acids (AA), alcohols (alc), alkaloids (alk), aromatics (aro), bioamines (bio), fatty acids and lipids (FAL), flavonoids (fla), organic acids (oAc), stillbenoids (stb) and terpenes (ter).

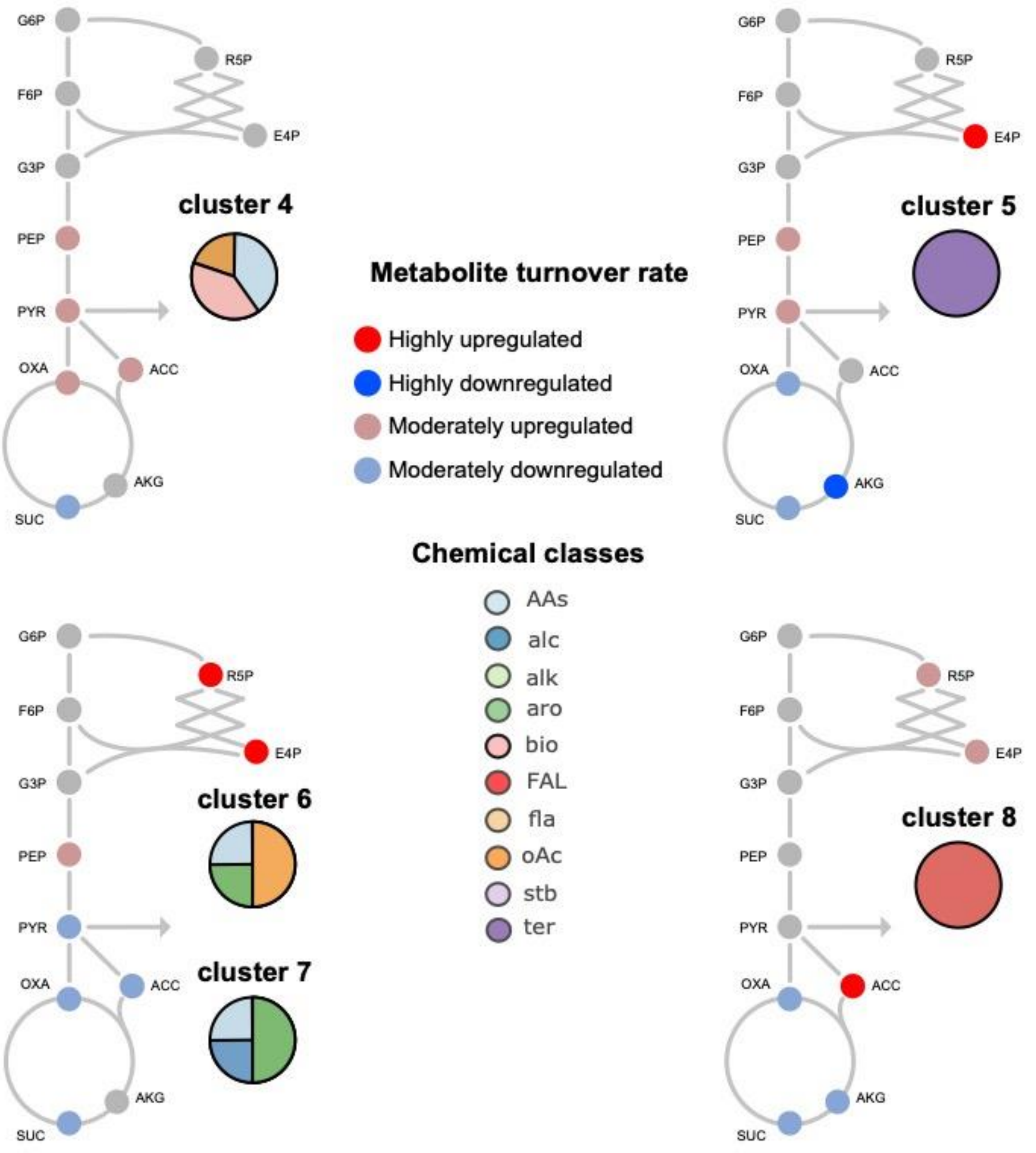


**Figure S7.-** Changes in precursors turnover required for optimal chemical production in yeast platform strains. Metabolites with a $|\log_2 FC| \geq 2$ in their turnover rate, when comparing a production scenario to wild-type simulations, are referred as "highly up/down-regulated" metabolic precursors. Metabolites with $1.5 < |\log_2 FC| < 2$ in their turnover rate are classified as moderately "up/down-regulated". The cluster classification is based on …?

### 3.- Supplementary Tables

**Table S1.** 102 chemicals simulated with ecYeastGEM.

| Name | Formulac | KEGG_ID | CHEBI_ID | MW | Origin | Chemical class | limitation |
|---|---|---|---|---|---|---|---|
| L-phenylacetylcarbinol | C9H10O2 | NA | 149766 | 150.174 | heterologous | alcohol | substrate |
| n-butanol | C4H10O | C06142 | 28885 | 74.23 | heterologous | alcohol | protein |
| docosanol | C22H46O | D03884 | 31000 | 326.61 | heterologous | alcohol | protein |
| (R,R)-2,3-butanediol | C4H10O2 | C03044 | 16982 | 90.121 | native | alcohol | substrate |
| 2-phenylethanol | C8H10O | C05853 | 49000 | 122.1644 | native | alcohol | substrate |
| glycerol | C3H8O3 | C00116 | 17754 | 92.09382 | native | alcohol | substrate |
| isoamylol | C5H12O | C07328 | 15837 | 88.14818 | native | alcohol | substrate |
| isobutanol | C4H10O | C14710 | 46645 | 74.1216 | native | alcohol | substrate |
| (S)-Reticuline | C19H23NO4 | C02105 | 16718 | 329.39 | heterologous | alkaloid | protein |
| psilocybin | C12H17N2O4P | C07576 | 8614 | 284.248101 | heterologous | alkaloid | protein |
| cinnamoyltropine | C17H22ClNO2 | NA | NA | 307.82 | heterologous | alkaloid | protein |
| choline | C5H14NO | C00114 | 15354 | 104.1708 | native | alkaloid | protein |
| hypoxanthine | C5H4N4O | C00262 | 17368 | 136.11162 | native | alkaloid | substrate |
| xanthine | C5H4N4O2 | C00385 | 17712 | 152.11102 | native | alkaloid | substrate |
| Nicotianamine | C12H21N3O6 | C05324 | 17721 | 303.3116 | heterologous | amino acid | protein |
| ergothioneine | C9H15N3O2S | C05570 | 4828 | 229.3 | heterologous | amino acid | protein |
| glutathione | C10H17N3O6S | C00051 | 16856 | 307.32 | native | amino acid | substrate |
| L-alanine | C3H7NO2 | C00041 | 16977 | 89.09322 | native | amino acid | substrate |
| L-arginine | C6H14N4O2 | C00062 | 16467 | 174.201 | native | amino acid | substrate |
| L-asparagine | C4H8N2O3 | C00152 | 17196 | 132.118001 | native | amino acid | substrate |
| L-aspartate | C4H7NO4 | C00049 | 17053 | 133.1027 | native | amino acid | substrate |
| L-citrulline | C6H13N3O3 | C00327 | 16349 | 175.18584 | native | amino acid | substrate |
| L-cysteine | C3H7NO2S | C00097 | 17561 | 121.158 | native | amino acid | substrate |
| L-glutamate | C5H8NO4 | C00025 | 29985 | 146.12136 | native | amino acid | substrate |
| L-glutamine | C5H10N2O3 | C00064 | 18050 | 146.14458 | native | amino acid | substrate |
| L-glycine | C2H5NO2 | C00037 | 15428 | 75.06664 | native | amino acid | substrate |
| L-histidine | C6H9N3O2 | C00135 | 15971 | 155.15468 | native | amino acid | substrate |
| L-homoserine | C4H9NO3 | C00263 | 15699 | 119.1192 | native | amino acid | substrate |
| L-isoleucine | C6H13NO2 | C00407 | 17191 | 131.175 | native | amino acid | substrate |
| L-leucine | C6H13NO2 | C00123 | 15603 | 131.17296 | native | amino acid | substrate |
| L-lysine | C6H15N2O2 | C00047 | 32551 | 147.19558 | native | amino acid | substrate |
| L-methionine | C5H11NO2S | C00073 | 16643 | 149.21238 | native | amino acid | substrate |
| L-phenylalanine | C9H11NO2 | C00079 | 17295 | 165.18918 | native | amino acid | substrate |
| L-proline | C5H9NO2 | C00148 | 17203 | 115.1305 | native | amino acid | substrate |

| | | | | | | | |
|---|---|---|---|---|---|---|---|
| L-serine | C3H7NO3 | C00065 | 17115 | 105.09262 | native | amino acid | substrate |
| L-threonine | C4H9NO3 | C00188 | 16857 | 119.1192 | native | amino acid | substrate |
| L-tryptophan | C11H12N2O2 | C00078 | 16828 | 204.22526 | native | amino acid | substrate |
| L-tyrosine | C9H11NO3 | C00082 | 17895 | 181.18858 | native | amino acid | substrate |
| L-valine | C5H11NO2 | C00183 | 16414 | 117.14638 | native | amino acid | substrate |
| ornithine | C5H13N2O2 | C01602 | 46912 | 133.169 | native | amino acid | substrate |
| 4-hydroxymandelic | C8H8O4 | C11527 | 16388 | 168.148 | heterologous | aromatic | substrate |
| cinnamate | C9H8O2 | C00423 | 15669 | 148.1586 | heterologous | aromatic | substrate |
| mandelic acid | C8H8O3 | C01984 | 17756 | 152.1473 | heterologous | aromatic | substrate |
| p-coumaric acid | C9H8O3 | C00811 | 32374 | 164.0473 | heterologous | aromatic | protein |
| rosmarinate | C18H16O8 | C01850 | 50371 | 360.31 | heterologous | aromatic | protein |
| Salidroside | C14H20O7 | C06046 | 9009 | 300.307 | heterologous | aromatic | protein |
| Tyrosol | C8H10O2 | C06044 | 1879 | 138.164 | heterologous | aromatic | protein |
| vanillin β-glucoside | C14H18O8 | NA | 179508 | 314.29 | heterologous | aromatic | protein |
| β-ionone | C13H20O | C12287 | 32325 | 192.3 | heterologous | aromatic | protein |
| putrescine | C4H14N2 | C00134 | 326268 | 90.1674 | native | bioamine | substrate |
| spermine | C10H30N4 | C00750 | 45725 | 206.372 | native | bioamine | substrate |
| ARA | C20H32O2 | C00219Ôø Ω | 15843 | 304.474 | heterologous | fatty acids and lipids | protein |
| DHA | C22H32O2 | C06429 | 28125 | 328.488 | heterologous | fatty acids and lipids | protein |
| EPA | C20H30O2 | C06428 | 28364 | 302.451 | heterologous | fatty acids and lipids | protein |
| Free fatty acids | NA | NA | NA | 396.64836 | native | fatty acids and lipids | substrate |
| laurate | C12H23O2 | C02679 | 18262 | 199.31 | native | fatty acids and lipids | protein |
| oleate | C18H34O2 | C00712 | 16196 | 282.4614 | native | fatty acids and lipids | substrate |
| palmitoleate | C16H29O2 | C08362 | 32372 | 253.40026 | native | fatty acids and lipids | substrate |
| triacylglycerol | C6H5O6R3 | C00422 | 17855 | 173.1003 | native | fatty acids and lipids | substrate |
| ergosterol | C28H44O | C01694 | 16933 | 396.64836 | native | fatty acids and lipids | substrate |
| catechin | C15H14O6 | C06562 | 15600 | 290.26 | heterologous | flavonoid | protein |
| genistein | C15H10O5 | C06563 | 28088 | 270.241 | heterologous | flavonoid | protein |
| kaempferol | C15H10O6 | C05903 | 28499 | 286.23 | heterologous | flavonoid | protein |
| naringenin | C15H12O5 | C00509 | 17846 | 272.257 | heterologous | flavonoid | protein |
| quercetin | C15H10O7 | C00389 | 16243 | 302.236 | heterologous | flavonoid | protein |
| Adipic acid | C6H10O5 | C023060 | 30832 | 146.14 | heterologous | organic acid | substrate |
| cis,cis-muconate | C6H6O4 | C02480 | 16508 | 142.11 | heterologous | organic acid | substrate |
| Itaconic acid | C5H6O4 | C00490 | 17240 | 130.1 | heterologous | organic acid | substrate |
| caffeic acid | C9H8O4 | C01197 | 36281 | 180.16 | heterologous | organic acid | protein |
| succinate | C4H6O4 | C00042 | 26806 | 118.09 | native | organic acid | substrate |
| (S)-lactate | C3H6O3 | C00186 | 16651 | 89.07 | native | organic acid | substrate |

| | | | | | | | |
|---|---|---|---|---|---|---|---|
| (S)-malate | C4H6O5 | C00149 | 15589 | 132.07156 | native | organic acid | substrate |
| 2-oxoglutarate | C5H6O5 | C00026 | 16810 | 144.08226 | native | organic acid | substrate |
| acetate | C2H4O2 | C00033 | 15366 | 60.052 | native | organic acid | substrate |
| citrate | C6H8O7 | C00158 | 16947 | 189.0997 | native | organic acid | substrate |
| fumarate | C4H4O4 | C00122 | 18012 | 116.0722 | native | organic acid | substrate |
| glycolate | C2H4O3 | C00160 | 17497 | 76.05136 | native | organic acid | substrate |
| hexanoate | C6H12O2 | C01585 | 17120 | 115.15034 | native | organic acid | protein |
| pyruvate | C3H3O3 | C00022 | 15361 | 87.05412 | native | organic acid | substrate |
| resveratrol | C14H12O3 | C03582 | 27881 | 228.25 | heterologous | stilbenoids | protein |
| astaxanthin | C40H52O4 | C08580 | 40968 | 596.841 | heterologous | terpene | protein |
| betaxanthin | C18H18N2O7 | C08565 | 8341 | 358.3 | heterologous | terpene | protein |
| β-carotene | C40H56 | C02094 | 17579 | 536.8726 | heterologous | terpene | protein |
| artemisinic acid | C15H22O2 | C20309 | 63749 | 234.33 | heterologous | terpene | substrate |
| alpha-Farnesene | C15H24 | C09665 | 10280 | 204.36 | heterologous | terpene | substrate |
| amorphadiene | C15H24 | C16028 | 52026 | 204.351 | heterologous | terpene | substrate |
| Geraniol | C10H18O | C01500 | 17447 | 154.25 | heterologous | terpene | substrate |
| Limonene | C10H16 | C06078 | 15384 | 136.24 | heterologous | terpene | substrate |
| linalool | C10H18O | C03985 | 17580 | 154.25 | heterologous | terpene | protein |
| lupeol | C30H50O | C08628 | 6570 | 426.72 | heterologous | terpene | protein |
| Oleanolic acid | C30H48O3 | C17148 | 37659 | 456.7 | heterologous | terpene | protein |
| patchoulol | C15H26O | C19755 | 7940 | 222.36 | heterologous | terpene | protein |
| protopanaxadiol | C30H52O3 | C20715 | 75950 | 460.73 | heterologous | terpene | protein |
| Santalene | C15H24 | C19736 | 10440 | 204.35 | heterologous | terpene | substrate |
| β-Amyrin | C30H50O | C08616 | 10352 | 426.72 | heterologous | terpene | substrate |
| glycyrrhetinic acid | C30H46O4 | C02283 | 30853 | 470.7 | heterologous | terpene | protein |
| lycopene | C40H56 | C05432 | 15948 | 536.873 | heterologous | terpene | protein |
| miltiradiene | C20H32 | C20711 | 65037 | 272.5 | heterologous | terpene | protein |
| nootkatone | C15H22O | C17914 | 81377 | 218.33 | heterologous | terpene | protein |
| squalene | C30H50 | C00751 | 15440 | 410.73 | heterologous | terpene | protein |
| taxadien-5alphayl-acetate | C22H34O2 | NA | 30042 | 330.5 | heterologous | terpene | protein |
| valencene | C15H24 | C17277 | 61700 | 204.35 | heterologous | terpene | protein |

**Table S4.**

Gene targets predicted for 2-phenylethanol increased production.

| Genes | Enzymes | Short names | Pathways | Target type | Enzyme type |
|---|---|---|---|---|---|
| YDR380W | Q06408 | ARO10 | sce00360 Phenylalanine metabolism | OE | Essential for production |
| YNL316C | P32452 | PHA2 | sce00400 Phenylalanine, tyrosine and tryptophan biosynthesis sce01110 Biosynthesis of secondary metabolites sce01130 Biosynthesis of antibiotics sce01230 Biosynthesis of amino acids | OE | Essential for production |
| YPR060C | P32178 | ARO7 | sce00400 Phenylalanine, tyrosine and tryptophan biosynthesis sce01110 Biosynthesis of secondary metabolites sce01130 Biosynthesis of antibiotics sce01230 Biosynthesis of amino acids | OE | Essential for production |
| YDR127W | P08566 | ARO1 | sce00400 Phenylalanine, tyrosine and tryptophan biosynthesis sce01110 Biosynthesis of secondary metabolites sce01130 Biosynthesis of antibiotics sce01230 Biosynthesis of amino acids | OE | Essential for production |
| YGL148W | P28777 | ARO2 | sce00400 Phenylalanine, tyrosine and tryptophan biosynthesis sce01110 Biosynthesis of secondary metabolites sce01130 Biosynthesis of antibiotics sce01230 Biosynthesis of amino acids | OE | Essential for production |
| YBR249C | P32449 | ARO4 | sce00400 Phenylalanine, tyrosine and tryptophan biosynthesis sce01110 Biosynthesis of secondary metabolites sce01130 Biosynthesis of antibiotics sce01230 Biosynthesis of amino acids | OE | Essential for production |

| | | | | | |
|---|---|---|---|---|---|
| YNL241C | P11412 | ZWF1 | sce00030 Pentose phosphate pathway sce00480 Glutathione metabolism sce01110 Biosynthesis of secondary metabolites sce01130 Biosynthesis of antibiotics sce01200 Carbon metabolism | OE | Essential for production |
| YHR163W | P38858 | SOL3 | sce00030 Pentose phosphate pathway sce01110 Biosynthesis of secondary metabolites sce01130 Biosynthesis of antibiotics sce01200 Carbon metabolism | OE | Optimal for production |
| YHR183W | P38720 | GND1 | sce00030 Pentose phosphate pathway sce00480 Glutathione metabolism sce01110 Biosynthesis of secondary metabolites sce01130 Biosynthesis of antibiotics sce01200 Carbon metabolism | OE | Optimal for production |
| YOR175C | Q08548 | ALE1 | sce00561 Glycerolipid metabolism sce00564 Glycerophospholipid metabolism sce01110 Biosynthesis of secondary metabolites | KD | Biomass coupled |
| YDR503C | Q04396 | LPP1 | sce00561 Glycerolipid metabolism sce00564 Glycerophospholipid metabolism sce01110 Biosynthesis of secondary metabolites | KD | Biomass coupled |
| YKL029C | P36013 | MAE1 | sce00620 Pyruvate metabolism sce01200 Carbon metabolism | KO | Unnecessary for production |
